# Comparative genomics of abiotic stress response in Norway spruce and Scots pine

**DOI:** 10.64898/2026.08.03.742458

**Authors:** Elena M van Zalen, Camilla Canovi, Vikash Kumar, Ellen Dimmen Chapple, David Castro, Sonja Viljamaa, Torgeir R Hvidsten, Vaughan Hurry, Nathaniel R Street

**Affiliations:** Umeå Plant Science Centre (UPSC), Department of Plant Physiology, Umeå University, 901 83 Umeå, Sweden; Faculty of Chemistry, Biotechnology and Food Science, Norwegian University of Life Sciences, 1433 Ås, Norway; Umeå Plant Science Centre (UPSC), Department of Forest Genetics and Plant Physiology, Swedish University of Agricultural Sciences, 901 83 Umeå, Sweden; Science for Life Laboratory, Department of Plant Physiology, Umeå University, 901 83 Umeå, Sweden

**Author notes:** Corresponding author: Nathaniel Street, Umeå Plant Science Centre (UPSC), Department of Plant Physiology, Umeå University, 901 83 Umeå, Sweden.

## Abstract

Norway spruce (*Picea abies*) and Scots pine (*Pinus sylvestris*) are dominant boreal forest species with globally important contributions to carbon capture and storage and sustain an extensive forestry industry. Both species have adapted to cold and episodic drought, yet each occupies a distinct ecological niche. How conserved their stress responses are, which features are lineage-specific, and how far mechanisms known from herbaceous angiosperms apply, remain open questions of importance in the face of ongoing climate change. We profiled roots and needles of both species under drought and cold, combining differential expression with an orthology-aware comparative co-expression framework that places each gene on a conservation–divergence gradient. Differential expression was largely organ- and stress-specific, yet cross-species overlap at the orthogroup level was extensive and increased with stress intensity. Comparative co-expression recovered a further conserved regulatory backbone that per-timepoint differential expression did not resolve, enriched for abscisic-acid-centred signalling, oxidative and osmotic-stress responses and growth suppression, and containing canonical stress response transcription-factor families including NAC, WRKY and bZIP/ABF. The breadth of co-expression conservation for a gene was coupled to purifying selection on its coding sequence and to network connectivity. Additionally, segmental duplicates shared between the species were enriched among conserved drought circuits, whereas lineage-specific duplicates were enriched among genes lacking conserved co-expression. Regulatory conservation and genome architecture thus describe a single conservation–divergence axis, providing an evolution-anchored criterion that helps separate candidate core regulators (which are conserved, network-central and constrained) from reactive change that differential expression alone cannot resolve.

**Significance statement:** Cold and drought are recurring threats to boreal forests, yet how conifers coordinate their responses, and how much of that response is evolutionarily conserved, has been difficult to establish. Comparing Norway spruce and Scots pine, two species separated by a deep evolutionary divergence, we show that the genes whose co-expression is most broadly conserved between the species are also those under the strongest purifying selection on their protein-coding sequences. As these two species diverged so long ago, this conserved regulatory core stands out as an evolutionary signal that simple comparisons of differentially expressed genes fail to capture. The approach provides an evolution-anchored way to distinguish genes central to the stress response from lineage-specific or reactive change.

## Introduction

The boreal forest is one of the largest terrestrial biomes and stores a substantial fraction of global terrestrial carbon (Pan et al. 2011; Bradshaw and Warkentin 2015). Norway spruce (*Picea abies* (L.) H. Karst.) and Scots pine (*Pinus sylvestris* L.) dominate the Fennoscandian boreal zone (Caudullo et al. 2016; Durrant et al. 2016) and underpin carbon balance with global importance and sustain a major forest economy (Kumar et al. 2021). In temperate and boreal regions both species are routinely exposed to cold and drought stress, which impose overlapping osmotic and oxidative effects and that elicit partially shared transcriptional responses (Kim et al. 2024). Climate change is expected to increase the frequency and severity of both stresses (Seidl et al. 2017; D’Orangeville et al. 2018), so resolving how Norway spruce and Scots pine respond is central to predicting forest resilience and breeding for tolerance.

Stress responses are assembled from deeply conserved signalling modules. Upon perceiving a stress cue, plants generate reactive oxygen species (ROS), accumulate abscisic acid (ABA) and mobilise Ca²⁺, triggering protein-kinase cascades that reprogramme transcription via conserved transcription factor (TF) families including CBF/DREB, NAC, WRKY, MYB, AP2/ERF and bZIP, although current knowledge derives mainly from herbaceous angiosperm models (Xu et al. 2022). Cold acclimation is broadly conserved, yet its timing and amplitude differ among species and tissues (Rihan et al. 2017; Preston and Sandve 2013). Non-freezing temperatures alter membrane fluidity and ionic homeostasis, elevate ROS and cytosolic Ca²⁺, and modulate hormone signalling (Örvar et al. 2000). In angiosperms the canonical cold axis, ICE1–CBF–COR, is driven by ICE1 activation of CBF/DREB regulons that induce cold-responsive genes (Chinnusamy et al. 2003; Chinnusamy et al. 2007). Homologues occur across plant lineages, but their roles are thought to differ in conifers. In Norway spruce an ICE1-like factor was identified as a central regulator of the cold response in both needles and roots, whereas the single *CBF1*/*CBF3*orthologue was only weakly and transiently cold-induced in roots and was unresponsive in needles (Vergara et al. 2022) and *CBFs* are not thought to be part of the conserved root cold response in conifers (Aro et al., in prep.). Drought stress induces the core ABA module (PYR/PYL/RCAR–PP2C–SnRK2) that drives stomatal closure and osmotic adjustment and, since freezing imposes cellular dehydration analogous to water deficit, cold and drought share much of this osmotic and protective machinery (Siminovitch and Cloutier 1983). Several early signals (ROS, Ca²⁺, ABA) are shared with drought, but downstream outcomes diverge under hydraulic constraints induced by water deficit.

Water availability is a primary determinant of growth and survival. Under drought, ABA accumulates in guard cells and drives stomatal closure, reducing transpiration, which protects the xylem but also restricts CO₂ uptake and growth, potentially precipitating carbon starvation (Cowan and Farquhar 1977; Pei et al. 2000; Hammond et al. 2019). Norway spruce and Scots pine both regulate water status conservatively, towards the isohydric end of the spectrum, limiting embolism (Irvine et al. 1998; Salmon et al. 2015; Walthert et al. 2024). What distinguishes them is water acquisition with the shallow plate-root system of Norway spruce confining uptake to surface soil that dries first, increasing hydraulic limitation under summer drought, whereas the deeper taproot of Scots pine buffers against surface drought and supports faster recovery (Brinkmann et al. 2018; Aldea et al. 2022).

The extent to which largely angiosperm-derived understanding of abiotic stress-response regulation applies to conifers is unclear and hard to test directly. Conifers and angiosperms diverged several hundred million years ago, and Norway spruce and Scots pine are themselves deeply diverged. In conifers, stress-response genes evolve at lower substitution rates than in angiosperms, though with comparable or somewhat elevated dN/dS, consistent with the slow gymnosperm molecular clock rather than uniformly stronger purifying selection (Buschiazzo et al. 2012; De La Torre et al. 2017; Garosi et al. 2025), and expression divergence between the two species tracks sequence evolution rather than positive selection (Hodgins et al. 2016). Identifying gene expression regulatory conservation or divergence using simple overlap of differentially expressed genes has limitations since regulatory divergence is pervasive even where coding-sequence orthology is conserved (Hartmann et al. 2022; Movahedi et al. 2012). Orthology-aware comparative co-expression instead tests whether the network-neighbourhood context of orthologues is conserved, giving greater power to recover conserved regulation that per-timepoint differential-expression comparison misses, and can help prioritise candidate core regulators over reactive, damage-associated change (Netotea et al. 2014; Proost and Mutwil 2018). This suits the deep *Picea–Pinus* divergence, where matched-timepoint expression comparison is confounded by when each gene crosses the significance threshold, while network-level conservation remains detectable.

In conifers it remains largely uncharacterised which genes are co-regulated under stress, whether their partnerships are conserved across the ∼140-My *Picea–Pinus* divergence (Yeaman et al. 2014), and how co-expression conservation relates to coding-sequence constraint, gene duplication and population-level variation. These questions are timely as pan-plant analyses show that orthogroups repeatedly recruited for local climate adaptation occupy central, broadly expressed positions in co-expression networks (Whiting et al. 2024), and that stress-response conservation decays with phylogenetic distance yet remains detectable across more than 700 My of divergence (Koh et al. 2026). Conifers are also not a uniform lineage with, for example, Norway spruce and Scots pine occupying different ecological niches and differing in their cold- and drought-response strategies (Caudullo et al. 2016; Durrant et al. 2016), making them well suited to separating a conserved core from lineage-specific regulation.

To address these questions, we profiled needles and roots of Norway spruce and Scots pine under controlled cold and drought, combining differential expression analysis with orthology-aware comparative co-expression, asking: (i) how much of the induced transcriptional response is shared between organs and species; (ii) whether comparative co-expression recovers a conserved regulatory core that differentially expressed gene (DEG) overlap misses because matched-timepoint comparison depends on when each gene crosses the significance threshold; and (iii) how the resulting spectrum of co-expression conservation relates to coding-sequence evolution, gene duplication and population-genomic variation. We found extensive organ- and species-specific variation overlying a conserved regulatory backbone, showing that the breadth of co-expression conservation is mirrored by coding-sequence constraint and genome architecture, and used this coupling to derive an evolution-anchored criterion that distinguishes candidate core regulators, conserved, constrained and network-central, from reactive change that differential expression analysis cannot resolve. We treat these as two readings of one axis: comparative co-expression places each gene on a conservation–divergence gradient, while molecular evolution and genome architecture reveal what separates the conserved core from the divergent periphery.

## Results

### Organ identity dominates the transcriptional response to cold and drought

Roots and needles of Norway spruce and Scots pine seedlings were sampled under cold and drought across a graded stress time course (Supplementary Figure S1). Drought was imposed as a progressive decline in soil water availability, sampled at mild and moderate stress, under severe drought at the onset of photosynthetic failure (as defined in Haas et al. 2021 Figure 1b and Figure S1 here), and after re-watering. Cold was applied as a stepwise reduction to 5 °C then −5 °C, sampling over ten days at each temperature. Principal component analyses (PCA) of the RNA-seq expression data were used to examine among-sample relationships within each dataset and to determine whether organs and species shared common response patterns. In both species, separation was strongest between organs rather than stress conditions with organ type accounting for the great majority of variance (∼85%; PC1), whereas stress conditions explained up to ∼12% (PC2) (Figure 1a–d). To quantify response magnitude, differentially expressed genes (DEGs) were identified. Drought triggered a strong root response at photosynthetic failure while needles had far fewer drought DEGs throughout, though their strongest response likewise occurred at photosynthetic failure. Under cold stress, conversely, needles showed a more pronounced transcriptional response than roots in both species, with DEG numbers generally increasing as treatment progressed. Gene Ontology (GO) enrichment analysis showed up-regulated genes enriched predominantly for defence and carbohydrate-metabolism processes (with water-deprivation responses among the drought-root genes), while down-regulated genes were enriched for signalling, secondary-metabolite biosynthesis and cell-wall/vascular processes (Supplementary Table S1).

**Figure 1.**
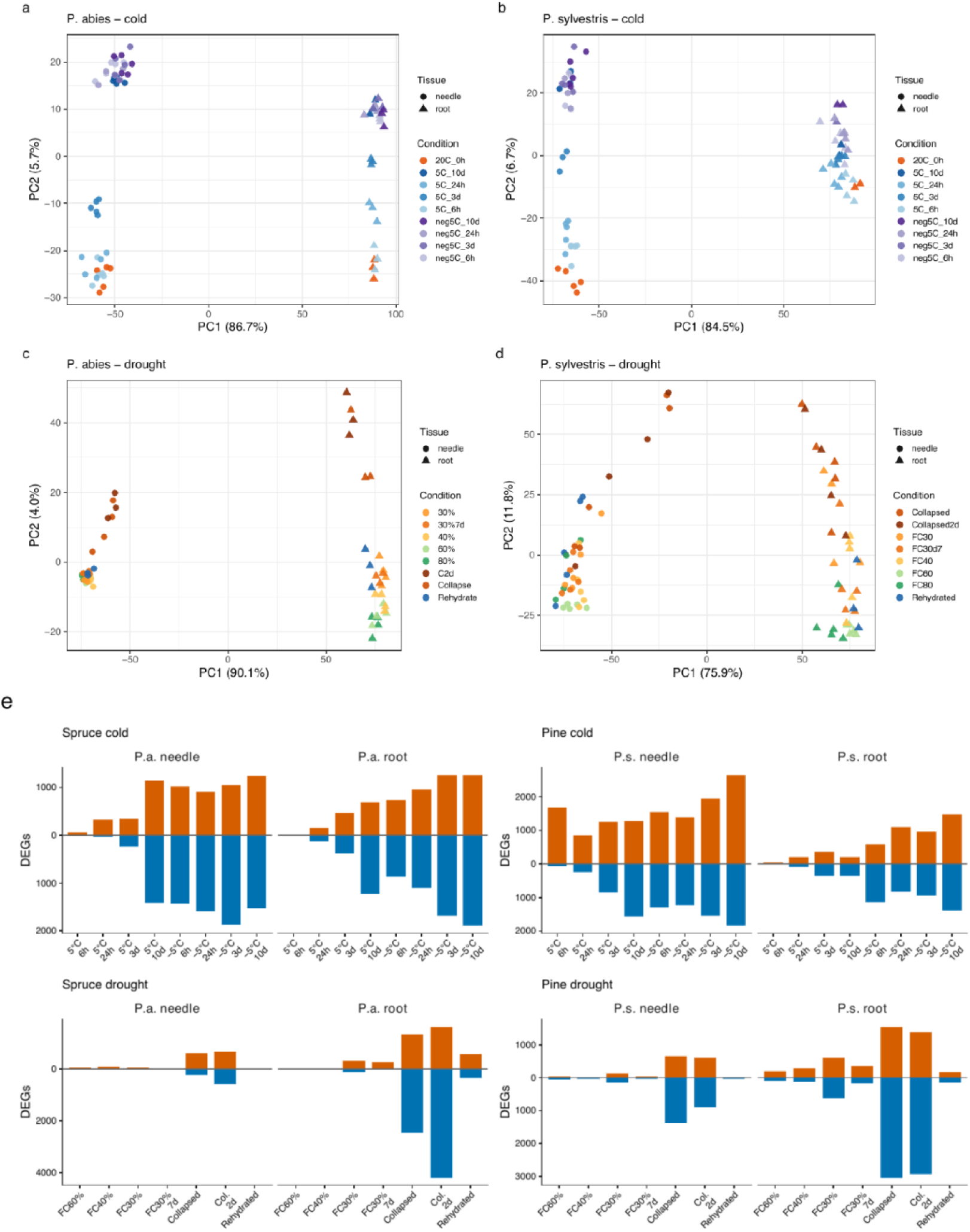
Transcriptional responses to cold and drought stress in Norway spruce and Scots pine. (a–d) Principal component analyses (PCA) of variance stabilizing-transformation (VST)-normalised RNA-seq expression data from *Picea abies* cold (a), *Pinus sylvestris* cold (b), *P. abies* drought (c) and *P. sylvestris* drought (d) experiments. Each point represents one biological replicate, coloured by experimental condition and shaped by tissue (circles = needles, triangles = roots). PC1 separates needle and root samples (76–90 % of the variance among the top 500 most variable genes); PC2 captures stress progression (4–12 %). (e) Stacked bar plots of differentially expressed genes (log2FC ≥ 2, *P*adj ≤ 0.01; red = up-regulated, blue = down-regulated) at each timepoint, arranged by species–stress–tissue. A vertical dashed line separates needle (left) from root (right) conditions within each panel.

### Abiotic stress largely activates organ- and stress-specific genes

We examined DEG overlap across datasets within each species (Figure 2a–b; full per-comparison DEG lists in Supplementary Table S2). In both Norway spruce and Scots pine, most DEGs were dataset-specific, the largest sets being drought-root, cold-needle and cold-root, indicating that each species primarily induced organ- and stress-specific transcriptional changes. Among shared genes, the cold-specific overlap (differentially expressed in both cold datasets but neither drought dataset) comprised 834 genes in Norway spruce and 1,303 in Scots pine while the drought-specific overlap was smaller, at 253 and 514 genes respectively. Within the cold-specific overlap, Norway spruce genes were enriched for signal transduction and glutathione and steroid metabolism, whereas Scots pine genes were enriched for photoprotective light responses (cellular responses to far-red light, UV-A and high light intensity). Within the drought-specific sets, Norway spruce genes were enriched for glutathione catabolism, response to gibberellin and regulation of defence response, and Scots pine genes for stomatal-complex patterning and formation and guard-mother-cell differentiation. Across both organs and stresses, 274 genes in Norway spruce and 344 in Scots pine were shared, representing a core set of candidate general stress-response genes common to cold and drought within each species. GO enrichment of this shared core recovered carbohydrate and energy-reserve metabolism in both species; galactose metabolism, glycogen biosynthesis and response to water in Norway spruce and galactose metabolism, glycogen biosynthesis and nitrate transport in Scots pine (P < 0.05, restricted to terms in the Arabidopsis (*Arabidopsis thaliana* (L.) Heynh.) GO universe) (Supplementary Table S1). These represent shared functional categories that may be conserved components of the responses to both stresses. Because these within-species sets were computed at the gene level against each species’ own annotation, they cannot reveal whether shared functions reflect orthologous responses, which the orthogroup-level comparisons below address.

**Figure 2.**
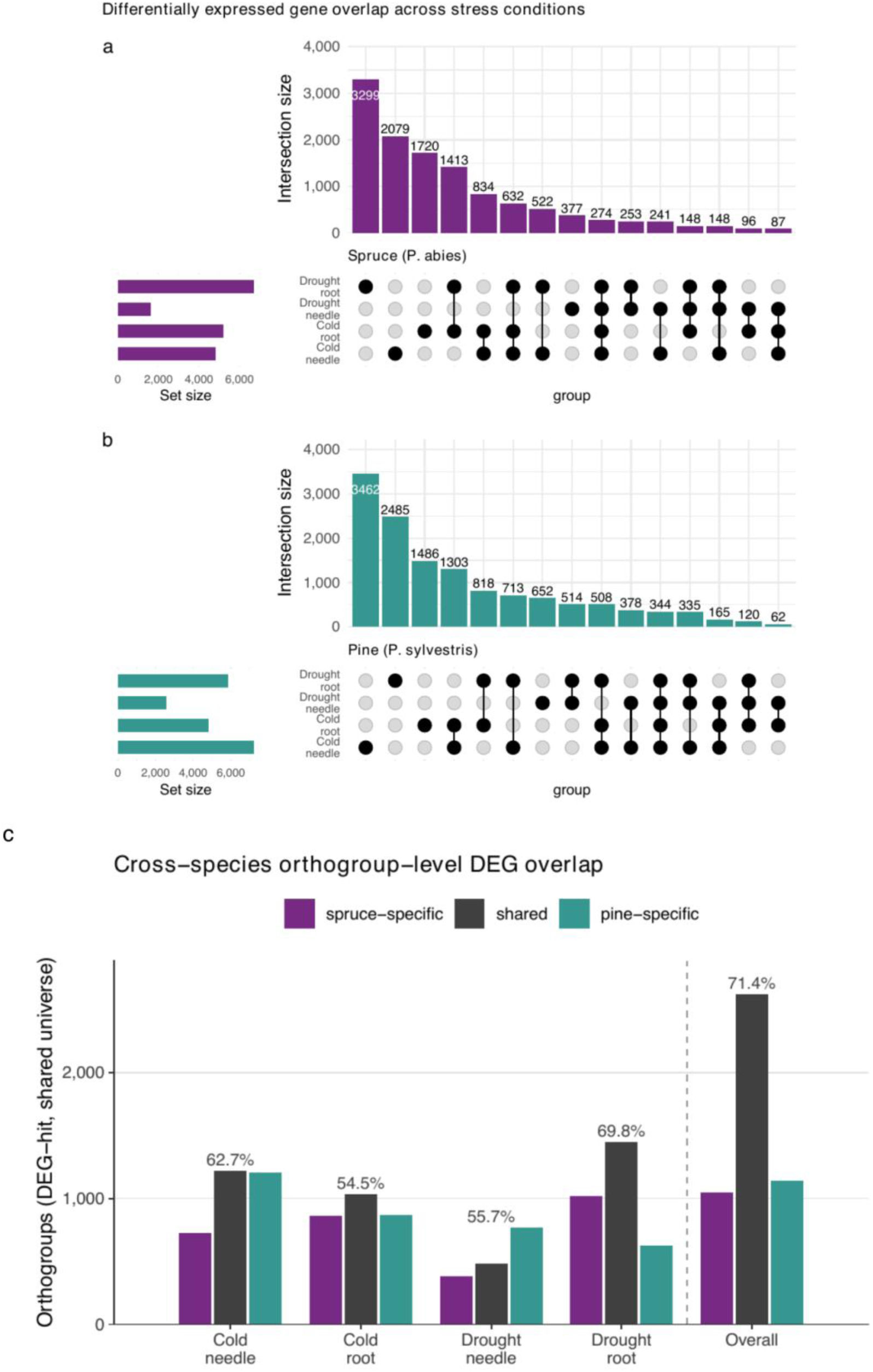
Overlap of differentially expressed genes within each species. UpSet plots of differentially expressed genes across the four stress–tissue datasets for (a) *Picea abies* and (b) *Pinus sylvestris*. Horizontal bars give total differentially expressed gene (DEG) counts per dataset; vertical bars give intersection sizes. The largest fractions are dataset-specific, with the largest shared set among cold-specific genes. (c) Cross-species comparison at the orthogroup level: for each matched sampling point (and pooled across all), the number of orthogroups differentially expressed only in *P. abies*, in both species (shared), or only in *P. sylvestris*, among the 11,943 orthogroups containing both a Norway spruce and a Scots pine gene. The percentage of the set of the smaller species that is shared is annotated above each group. Percentages and counts are computed among the 11,943 orthogroups containing a gene in both species; differentially expressed genes in single-species orthogroups (8.5% of *P. abies* and 10.9% of *P. sylvestris* DEG orthogroups) have no counterpart in the other species and are necessarily excluded.

To compare the two species on a common footing, we mapped the DEGs of each species to orthogroups and asked, at each matched sampling point, how many orthogroups contained DEGs in both species (Figure 2c; universe of 11,943 orthogroups containing both a Norway spruce and a Scots pine gene). An orthogroup was counted as shared when it contained at least one differentially expressed gene from each species, a gene-set criterion at the orthogroup level that does not require one-to-one orthology and allows an orthogroup to contain several genes from either species. When considering all sampling points together for each stress, orthogroup-level overlap between the species was substantial: across stresses and organs, 2,625 orthogroups were differentially expressed in both species, 71% of the DEG orthogroups of the species with the smaller set and 2.3-fold more than expected by chance (hypergeometric P < 10⁻³⁰⁰). Where both species responded, they did so predominantly in the same direction: of the 1,646 shared orthogroups with an unambiguous direction in both species, 85.6% were directionally concordant, far exceeding the concordance expected if response directions were independent given the marginal up/down bias in each species (binomial P = 1.3 × 10⁻¹⁴¹), and concordant down-regulation outnumbered concordant up-regulation by roughly 2.5-fold (1,004 versus 405 orthogroups). Furthermore, overlap was significant at every sampling point (55–70% of the set of the smaller species shared per point; 3.1–5.3-fold enrichment; hypergeometric P < 10⁻²⁰⁰ at every point, largest P = 7.9 × 10⁻²⁷¹ at drought-needle).

The size of the shared fraction nonetheless varied along the stress progression. At early, mild timepoints few genes were differentially expressed in either species and cross-species overlap was minimal (for example 6.1–17.6% of cold-root DEG orthogroups at 5 °C, 6 h), whereas the shared orthogroup response accumulated with stress intensity, reaching roughly 58–76% of the DEG orthogroups of each species at the most severe cold and drought-root timepoints (for example 67% of Norway spruce and 62% of Scots pine cold-needle DEG orthogroups, and 69% and 76% of drought-root DEG orthogroups). The drought-needle comparison remained more asymmetric (63% of Norway spruce but 37% of Scots pine DEGs lay in shared orthogroups at photosynthetic failure) (Figure S2; Supplementary Table S3). This stress-intensity dependence indicates that overlap measured at any single matched timepoint understates the conserved response, because a shared, responding orthogroup is counted only when both species happen to cross the significance threshold at the same sampling point. For drought in particular, the response under the conditions applied was concentrated into a sharp transition at photosynthetic failure rather than spread evenly across the time-course, so the sampling point at which a gene is declared differentially expressed is especially sensitive. A genuine divergence in response timing was reported for boreal-tree roots across a wider species panel (Aro et al., in prep.); here, between these two conifers, the reduced matched-timepoint overlap is more parsimoniously attributed to this threshold-crossing effect than to divergent timing.

### Comparative co-expression reveals a conserved regulatory backbone

DEG analysis of matching timepoint sharply reduced the shared fraction of responsive genes relative to pooling all timepoints (from 58–76% of each species’ DEG orthogroups at the most severe timepoints down to as little as 6.1–17.6% at the mildest), indicating that a shared, responding orthogroup is registered only when both species cross the significance threshold at the same matched point. To remove this dependence on matched-timepoint calling, we next used comparative co-expression analysis to identify orthologs with shared co-expression neighbourhoods (co-expressologs). Applied across orthogroups, a comparative co-expression analysis recovered a shared regulatory component beyond that captured by DEG overlap: across the four pairwise stress–tissue comparisons, a set of orthogroups retained significant co-expressologs in all four contexts (Figure 3a), defining a conserved co-expression backbone. This set contained TFs central to stress regulation with co-expressologs conserved across all four comparisons including NAC, WRKY, MYB, AP2/ERF, bZIP/ABF, HSF, LBD and GLK families (Figure 3b). Differentially expressed TF co-expressolog pairs partitioned clearly between cold- and drought-responsive sets (Figure 3c).

**Figure 3.**
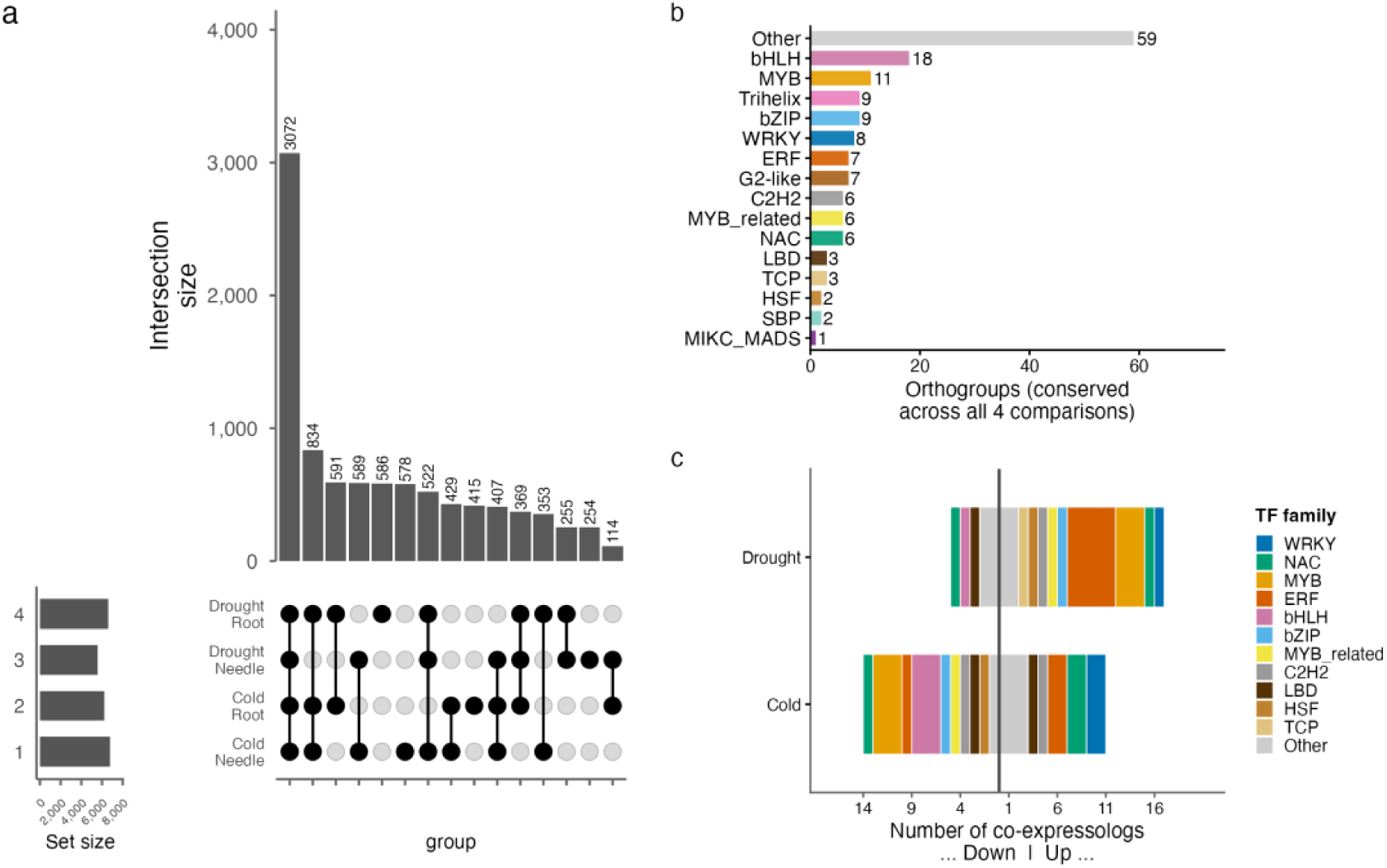
Cross-species co-expressolog conservation and transcription-factor composition. (a) UpSet plot of orthogroups with at least one significant co-expressolog (MaxpVal < 0.01) in each of the four pairwise stress–tissue comparisons; intersections of ≥ 10 orthogroups are shown. (b) Transcription factor (TF) families represented among orthogroups with co-expressologs conserved across all four pairwise comparisons (PlantTFDB annotation of *Picea abies* gene models; 1,954 TFs annotated); bars show orthogroup count per family. (c) TF co-expressolog pairs differentially expressed under drought (bottom) or cold (top) stress (DroughtSum ≥ 60 and ColdSum ≥ 40, respectively; differentially expressed in both species). Horizontal stacked bars show counts of up-regulated (right) and down-regulated (left) TF pairs, coloured by TF family; axis values are absolute counts. Unlike Figure 2 (DEG overlap within each species), this UpSet plot summarises orthogroups that share conserved cross-species co-expressologs across the four pairwise comparisons.

To test whether co-expressologs shared expression trajectories or encompassed shifted or inverted stress responses, we calculated their expression correlation across the stress time-course.Expression profiles were strongly conserved between species with a median Norway spruce–Scots pine correlation of 0.585 in needles and 0.786 in roots under drought (86% and 97% of orthogroups positively correlated), and 0.704 in needles and 0.634 in roots under cold (93% and 85% positive). These greatly exceeded a shuffled-partner null (median correlation 0.103 for drought and 0.046 for cold; Wilcoxon *P* < 2.2 × 10⁻¹⁶ for both stresses; Figure 4b). Conserved co-expressologs therefore not only exist but track one another quantitatively through the stress response in both species (Figure 4c,d). The reduced matched-timepoint DEG overlap we observed thus reflects threshold effects inherent to DEG-based comparisons rather than time-shifted stress responses.

Classifying orthogroups by the direction of their response separated a smaller up-regulated set from a larger down-regulated set under drought (312 up, 772 down) and a more balanced split under cold (407 up, 438 down). GO enrichment of these direction groups (Supplementary Table S4; plant-relevant terms) recovered a conserved stress trade-off. Up-regulated conserved co-expressologs were enriched for abscisic-acid-centred and protective metabolism including response to abscisic acid, trehalose and xanthophyll biosynthesis, and regulation of auxin biosynthesis and starch metabolism under drought, and wounding response, iron-sulfur cluster assembly and lipid-droplet organisation under cold. Down-regulated orthogroups were enriched for growth and cell-proliferation processes including carbohydrate metabolism, DNA-replication initiation, cell-wall modification, chlorophyll biosynthesis and procambium development. The conserved (shared) co-expression response thus reflected a common reallocation from growth to abscisic-acid-driven protection in both conifers.

**Figure 4.**
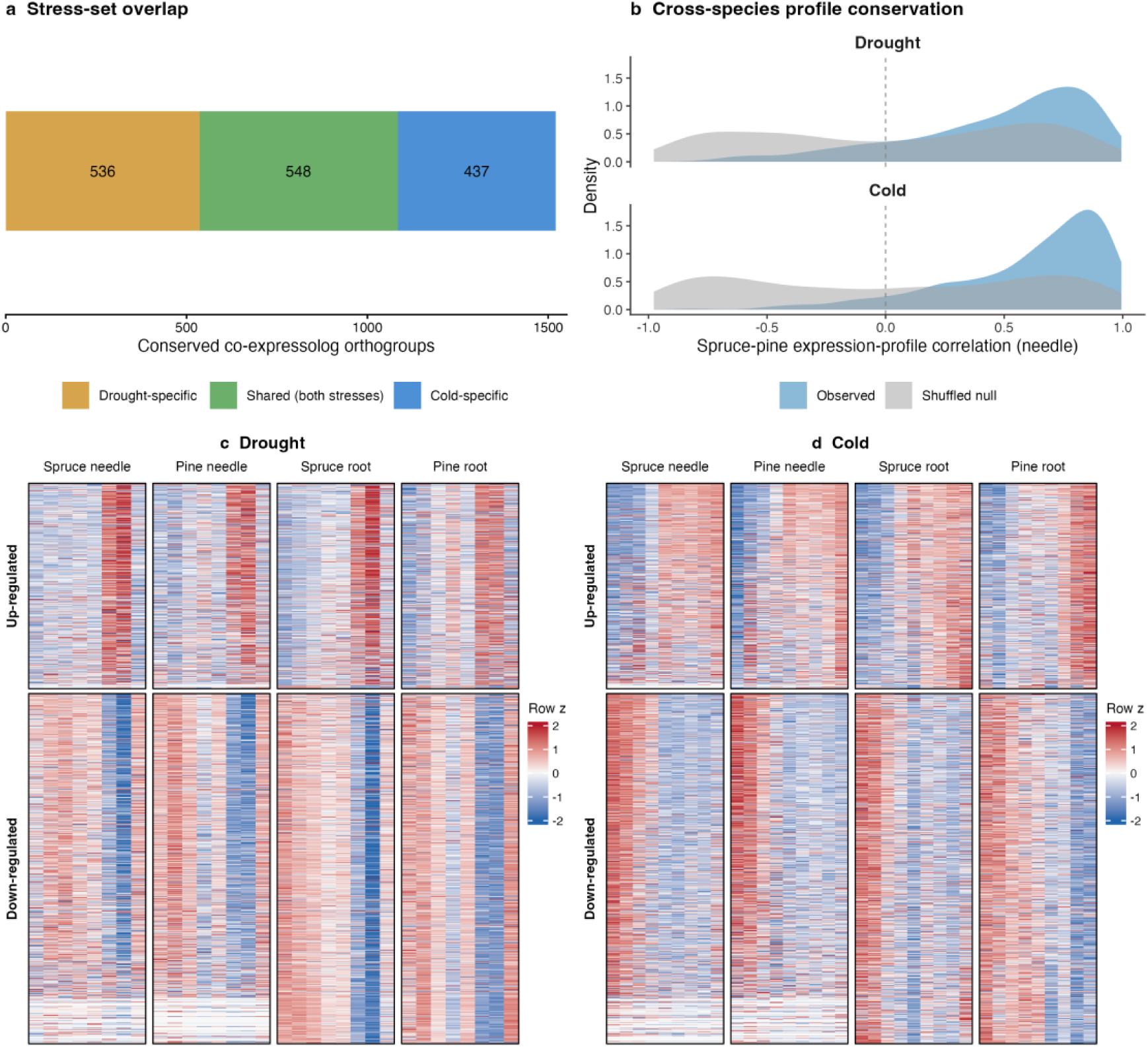
Conserved co-expressologs show conserved expression dynamics during stress. (a) Overlap of the drought- and cold-responsive conserved co-expressolog orthogroup sets (differentially expressed in both species; DroughtSum ≥ 60 / ColdSum ≥ 40). (b) Cross-species conservation of expression profile, shown as the density of the Pearson correlation between the *Picea abies* and *Pinus sylvestris* median expression trajectories (needle) for the observed orthogroup pairs versus a shuffled-partner null. (c, d) Heatmaps of median expression for (c) drought (1,084 orthogroups) and (d) cold (985 orthogroups) conserved co-expressologs; each row is one orthogroup and columns are the stress stages within four blocks (Norway spruce needle | Scots pine needle | Norway spruce root | Scots pine root), z-scored within each block, with rows split by response direction (up- versus down-regulated).

### Coding-sequence constraint tracks the breadth of co-expression conservation

Given that co-expression conservation defined a regulatory conservation–divergence axis spanning conserved in all comparisons to conserved in none, we asked whether this axis is reflected in coding-sequence evolution. The expectation tested was that genes with conserved stress co-regulation between the two species should be most constrained in coding sequence (low ratio of non-synonymous to synonymous substitution, dN/dS), whereas genes that have lost conserved co-regulation should be freer to diverge. To this end, we classified genes into five categories by the extent of cross-species co-expression across the four stress comparisons: conserved (co-expressolog pairs in all four; n = 1,615 Norway spruce and 1,621 Scots pine genes), cold-specific (n = 2,112 and 2,136, respectively), drought-specific (n = 1,742 and 1,745, respectively), multi-tissue (pairs in two or three comparisons spanning both stresses; n = 4,635 and 4,659, respectively) and not co-expressed (not_coex; no significant cross-species co-expression; n = 3,035 and 3,063, respectively). GO enrichment against the conserved co-expression universe revealed distinct functional signatures: conserved co-expressologs were enriched for core nucleic-acid-processing functions, including endonuclease activity (P = 1.0 × 10⁻³⁰), zinc ion binding (P = 1.0 × 10⁻³⁰) and RNA binding (P = 7.9 × 10⁻²⁸); cold-specific ones for RNA modification (P = 1.9 × 10⁻⁷) and RNA binding (P = 8.4 × 10⁻⁷); drought-specific ones for nuclear components (nucleus, P = 3.3 × 10⁻⁵; chromosome, P = 1.9 × 10⁻⁴). Multi-tissue co-expressologs showed the strongest overall enrichment, dominated by nucleic-acid binding and RNA processing (zinc ion binding, P = 7.8 × 10⁻²⁷; RNA binding, P = 2.8 × 10⁻²⁵; RNA modification, P = 9.9 × 10⁻¹⁷). Not_coex genes were most enriched for nucleoplasm (P = 8.9 × 10⁻¹⁰) and DNA binding (P = 1.2 × 10⁻⁷). Full enrichment results for all five categories are given in Supplementary Table S5.

There was a monotonic decrease in dN/dS with breadth of co-expression conservation across categories. Using codon-aware maximum-likelihood estimates (Yang and Nielsen 2000) for 13,519 cross-species orthologue pairs with usable dN/dS (of 13,523 pairs on the backbone), median dN/dS fell from not_coex (0.355) through cold-specific (0.317), drought-specific (0.316) and multi-tissue (0.308) to conserved (0.287). Treated as a continuous variable, dN/dS declined significantly with conservation breadth (Spearman ρ = −0.176, P = 5.2 × 10⁻⁹⁵; Figure 5a; Supplementary Table S6). The reciprocal analysis of the Scots pine co-expression axis showed the same decline (Spearman ρ = −0.19, P = 1.9 × 10⁻¹⁰⁶; Figure 5b). We defined the dN/dS backbone at the pairwise level, with each pair containing exactly one Norway spruce and one Scots pine gene, and with pairwise orthogroups holding more than one gene from either species excluded rather than being reduced to a representative pair. It should be noted that this pairwise criterion does not require single-copy status in the wider gene family. Indeed, of the 13,519 pairs with a usable dN/dS value, 11,166 (82.6%) belonged to hierarchical orthogroups (N10 level) that were single-copy, while the remaining 17.4% were pairwise 1:1 between Norway spruce and Scots pine but were nested within hierarchical orthogroups carrying additional paralogues elsewhere. Restricting the analysis to the 11,166 fully single-copy pairs (i.e. single copy in all species included in the orthology analysis) recovered essentially the same relationship (Spearman ρ = −0.166, P = 2.1 × 10⁻⁶⁹), confirming the gradient reflects coding-sequence constraint rather than an artefact of dN/dS estimation in multi-copy families. Genes with the broadest cross-species stress co-expression conservation are thus also under the strongest purifying selection on their coding sequences. To compare this gradient to a developmental context, we overlaid our cross-species dN/dS estimates onto the wood-formation co-expression cliques of Rodríguez et al. (2026), which classified genes by co-expression conservation across six species. Norway spruce genes in cliques conserved across all six species showed significantly lower dN/dS than those in lineage-differentiated or gymnosperm-specific cliques (median 0.262 versus 0.303 and 0.283; Kruskal–Wallis P = 0.004; conserved versus gymnosperm-specific P = 0.020 and versus differentiated P = 0.002), indicating that the coupling between coding-sequence constraint and co-expression conservation is a general property of conserved co-regulation, not specific to stress, with lineage-differentiated and gymnosperm-specific genes correspondingly less constrained.

**Figure 5.**
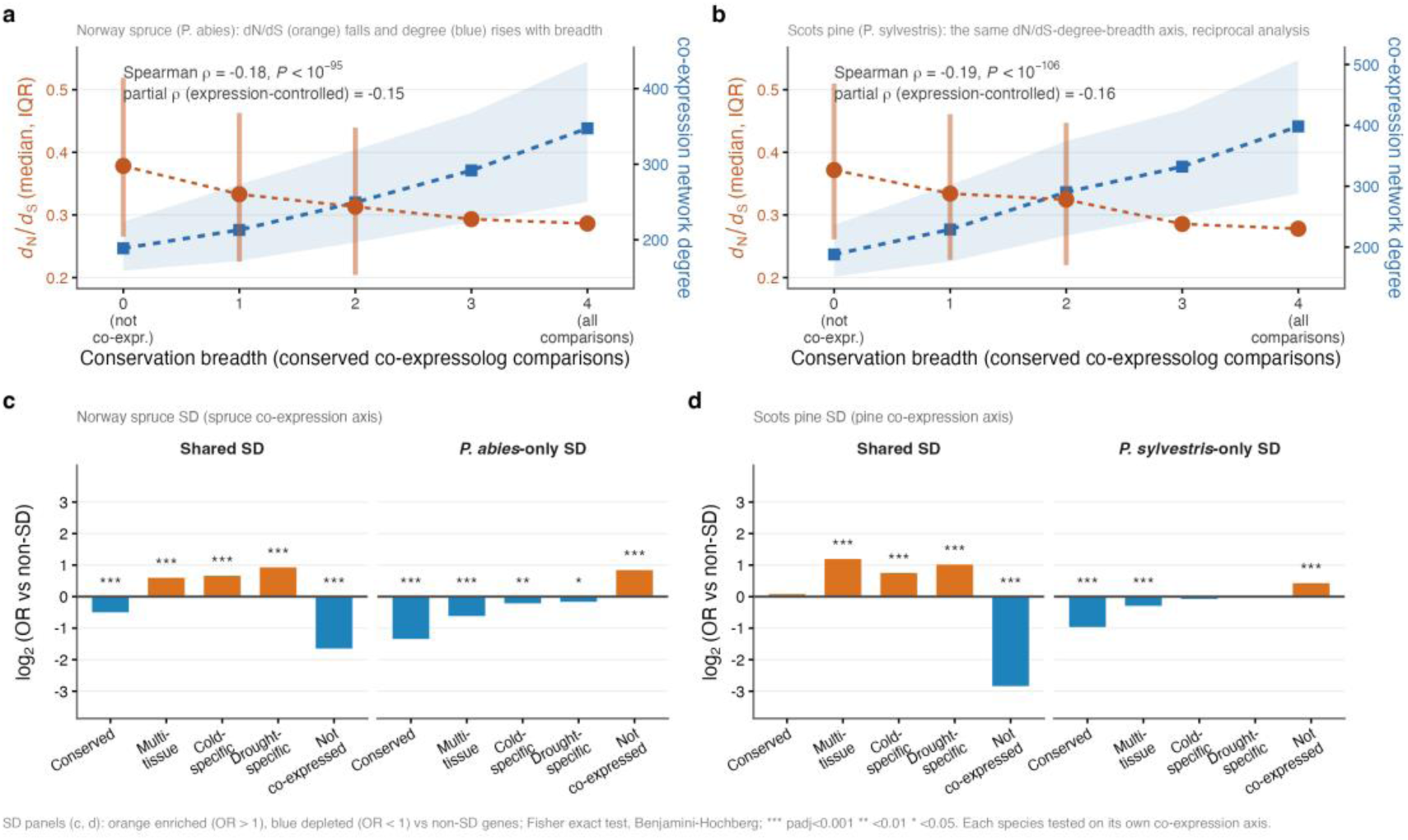
Evolutionary and genomic features of co-expression conservation categories. (a) dN/dS (non-synonymous:synonymous substitution ratio, estimated under the Yang–Nielsen (2000) maximum-likelihood model) for 13,519 cross-species orthologue pairs with usable dN/dS (of 13,523 pairs on the backbone) plotted against conservation breadth, the number of the four stress–tissue comparisons in which the orthogroup of a gene retains a significant conserved co-expressolog (0–4). Orange: median dN/dS (points) and interquartile range (bars), with the dashed line connecting breadth-level medians; blue (right axis): median co-expression network degree with interquartile range. Annotated statistics are the Spearman correlation of dN/dS with conservation breadth (Norway spruce ρ = −0.18) and the partial correlation controlling for expression level. (b) The reciprocal analysis on the Scots pine co-expression axis, showing the same dN/dS decline and network-degree increase with conservation breadth (Scots pine ρ = −0.19). (c, d) Segmental-duplicate (SD) gene enrichment across the five co-expression categories, shown reciprocally for Norway spruce (c) and Scots pine (d): in each species, shared_SD (independently duplicated in both species) and lineage-specific SD (*Picea abies*-only in c, *Pinus sylvestris*-only in d), tested on that species’ own co-expression axis. Bars show log2 (odds ratio) from Fisher’s exact test against the non-SD background; orange = enriched (OR > 1), blue = depleted (OR < 1); all bars Padj < 0.05 (Benjamini–Hochberg correction).

Two correlated gene properties could generate the above gradient without any specific link between co-expression conservation and coding-sequence constraint: broadly conserved co-expressologs are more central in the co-expression network (median network degree 374 in the conserved category versus 195 in not_coex) and tend to be more highly expressed, and both higher connectivity and higher expression are themselves linked to stronger purifying selection (Mähler et al. 2017). Controlling jointly for network degree and expression attenuated but did not remove the association between conservation breadth and dN/dS (partial Spearman ρ = −0.135, P = 1.4 × 10⁻⁵³; raw ρ = −0.152 on the same genes), consistent with connectivity, expression and coding-sequence constraint being coupled facets of a single conservation axis. The association is attenuated because these facets are correlated, yet constraint retains an independent link to conservation breadth beyond connectivity and expression. An independent, within-species measure of selection further supported this with median pN/pS, estimated from Norway spruce population resequencing (Ahlgren Kalman et al. 2025) increasing from 0.265 in the conserved category to 0.283 in not_coex (Kruskal–Wallis P = 3.3 × 10⁻⁸; conserved versus not_coex P = 4.2 × 10⁻⁶) This confirms that the constraint gradient is not only an artefact of between-species dN/dS estimation.

### Duplication and structural variation shape the divergent layer

The conifer genome is notable for its high content of segmental duplications (SDs), a potential engine of evolutionary novelty (Ahlgren Kalman et al. 2025). We therefore asked how duplicated genes are distributed across conserved versus lineage-specific layers of the cold and drought stress response. Segmental duplicate (SD) gene classes showed contrasting enrichment across expression conservation categories (Figure 5b). Genes duplicated independently in both Norway spruce and Scots pine (shared_SD; n = 1,874) were strongly enriched in the drought-specific category (OR = 1.93; Padj = 4.3 × 10⁻²⁶), more weakly in the cold-specific category (OR = 1.60; Padj = 2.4 × 10⁻¹⁴), and markedly depleted from the not_coex category (OR = 0.32; Padj = 3.2 × 10⁻⁹⁵) relative to non-SD genes. This indicates that independently duplicated genes are over-represented in conserved drought-specific co-regulatory circuits and that the same gene families were independently retained as duplicates in both lineages and channelled into conserved drought circuits, suggesting shared selective maintenance of these duplicated drought-responsive genes; both copies remained co-expressed in the network in 74% of shared-duplicate pairs. By contrast, lineage-specific duplicates were enriched in the not_coex category in both species: Norway spruce-specific duplicates (spruce_only_SD; n = 3,186; OR = 1.82; Padj = 8.5 × 10⁻⁵⁸) and, in a reciprocal analysis on the Scots pine co-expression axis, Scots pine-specific duplicates (pine_only_SD; n = 3,299; OR = 1.36; Padj = 2.6 × 10⁻¹⁷). In both species shared_SD duplicates were instead depleted from not_coex (Norway spruce OR = 0.32; Scots pine OR = 0.14) and enriched among the co-expressolog categories. This symmetry indicates that the partitioning of duplicated genes by co-expression conservation, shared duplicates into the conserved backbone and lineage-specific duplicates into the non-conserved layer, is a shared property of both lineages rather than an artefact of anchoring the analysis on one species.

### Mechanisms of species-specific regulation

To characterise the genes driving species-specific stress responses, we specifically asked why a large number of stress-responsive Norway spruce genes (1,983) lacked a conserved cross-species co-expressolog in Scots pine, assigning each to one of four mechanisms (Supplementary Figure S3a): diverged co-expression of a shared 1:1 orthologue; no 1:1 Scots pine orthologue, either no orthologue at all or membership of an expanded gene family; a 1:1 orthologue that was not stress-responsive; or a 1:1 orthologue not expressed under stress. Strikingly, very few of these genes (38 genes, 1.9%) were stress-responsive in both species but had diverged in co-expression. The species-specific response was instead dominated by the absence of a 1:1 orthologue: 1,859 genes (93.7%) had no 1:1 Scots pine orthologue at all. The remaining genes had a 1:1 orthologue that was either not stress-responsive (84 genes, 4.2%) or not expressed under stress (2 genes, 0.1%).

Species-specific stress responses in these conifers are therefore written largely into gene content, lineage-specific gene gain and gene-family expansion, rather than into the regulation of shared single-copy genes. Consistent with an origin in gene-family evolution, GO enrichment of the 1,859 lineage-specific genes was strongest for signal transduction (P = 1.1 × 10⁻²³) and protein phosphorylation (P = 1.6 × 10⁻⁹), indicating expansion of kinase-signalling families. The same architecture held in Scots pine with 2,809 ostress-responsive not_coex genes, of which 95.3% likewise lacked a 1:1 Norway spruce orthologue and only 1.2% reflected diverged regulation, and with the same GO signature (signal transduction P = 5.5 × 10⁻²¹; protein phosphorylation P = 1.6 × 10⁻³). Lineage-specific expansion of signalling-gene families is thus a shared, parallel route to regulatory divergence in both conifers.

Population-level positive selection signals, available only for Norway spruce, reinforced the above stratification. Genes with the strongest selection signatures, detected by both XP-EHH (north–south contrasting selection) and iHS (within-population sweep; Ahlgren Kalman et al. 2025), were enriched in the cold-specific category (OR = 1.24; padj = 0.0018) and significantly depleted from the conserved category (OR = 0.65; padj = 4.7 × 10⁻⁷), indicating that recent positive selection preferentially targets the cold-specific regulatory layer while avoiding the broadly conserved stress-response core. These genes occupied the network periphery (n = 2,712; median co-expressolog degree = 1, versus 3 for background genes; Wilcoxon P = 3.1 × 10⁻²³²) and showed roughly 5-fold higher median expression (median baseMean 321 versus 61 for background; Wilcoxon P < 1 × 10⁻³⁰⁸), suggesting that selection on this cold-specific layer targets specific, highly expressed effector genes rather than network hubs. Consistent with this climate-axis specificity, 12 temperature-associated GWAS genes (bio10 and bio11; Ahlgren Kalman et al. 2025) were differentially expressed under cold stress (5 up-regulated, 7 down-regulated), directly linking genetically encoded climate adaptation to the transcriptional responses documented here.

A *CHS3* segmental-duplicate pair on chromosome 9 illustrates this divergence at single-gene resolution (Figure 6). The two copies, which arose by a recent duplication, have diverged sharply in expression with one copy (PA_chr09_G004115) being induced specifically in cold-needles while the other (PA_chr09_G004116) is broadly expressed, despite strong purifying selection on both coding sequences (dN/dS = 0.261). Both copies are additionally subject to presence–absence variation (PAV) in the Norway spruce standing population, being absent or nearly so across the southern range (present in 0 of 26 and 2 of 26 southern individuals, respectively, versus all 26 northern individuals).

**Figure 6.**
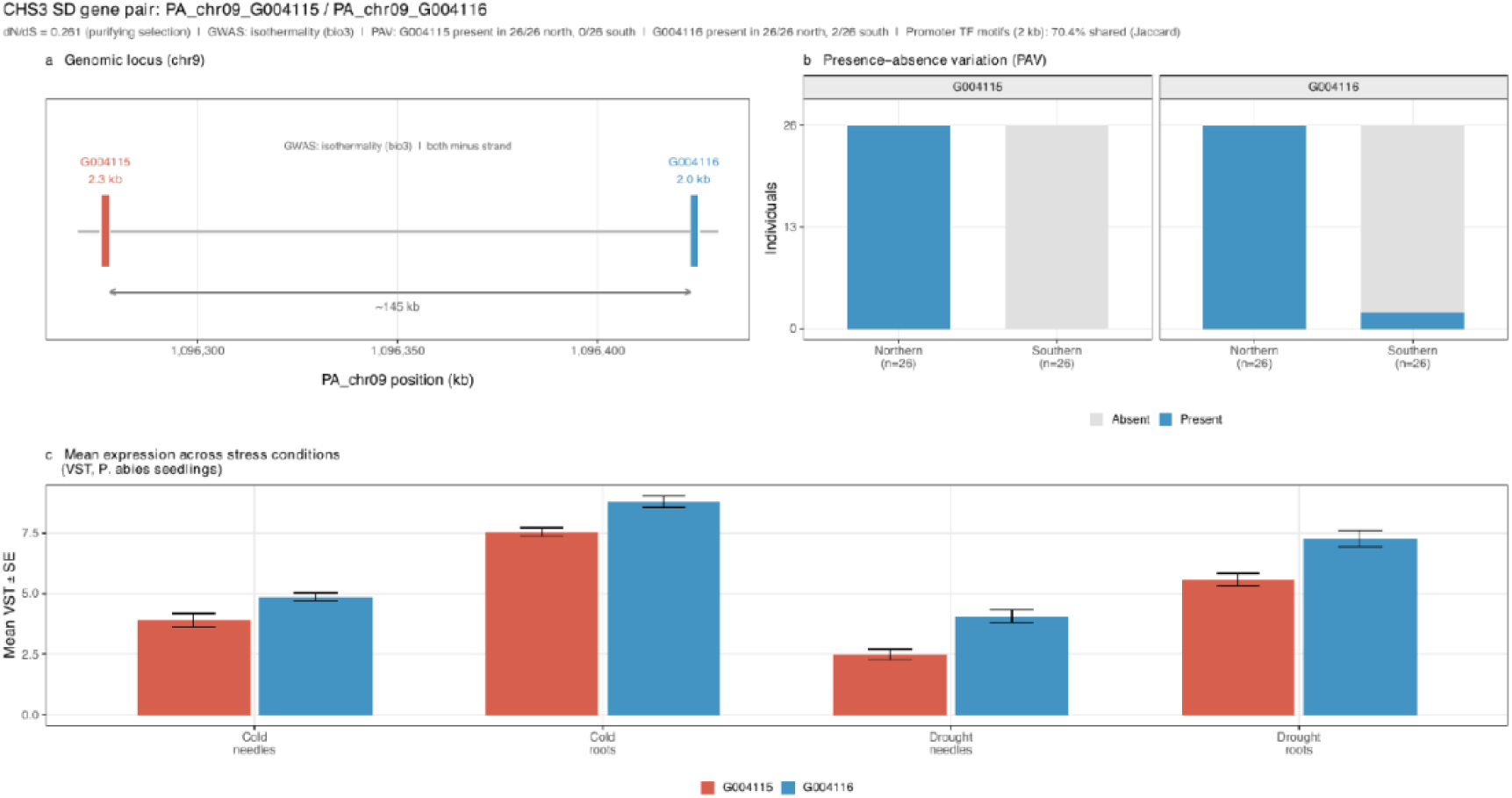
Case study: expression divergence of the *CHS3* segmental-duplicate pair (PA_chr09_G004115 and PA_chr09_G004116). (a) Genomic arrangement on *Picea abies* chromosome 9 (minus strand); positions are relative to the G004115 transcription start site. The paralog-pair age (Ks, synonymous distance) and coding-sequence selection (dN/dS) are annotated above. (b) Presence–absence variation (PAV) across northern and southern *P. abies* population panels (n = 26 individuals each). (c) Mean VST-normalised expression (± SEM across biological replicates) of each gene across the four *P. abies* stress datasets; G004115 displays cold-needle-specific expression, G004116 is broadly expressed. The chromosome-9 chalcone synthase gene family, in which this pair is nested, is shown as a maximum-likelihood phylogeny in Supplementary Figure S4.

The *CHS3* single-gene observation prompted us to ask whether population-level structural instability marks the divergent layer more generally. Across the standing-population PAV set, genes lacking conserved cross-species co-expression (not_coex) showed a weak but significant enrichment for PAV (OR = 1.77, P = 0.006; Benjamini–Hochberg-adjusted P =0.032 across the five co-expression categories; n = 95 PAV genes; Supplementary Figure S3b), with no significant enrichment in any other category. Given the modest effect size and the single standing-population dataset available, we treat this as an exploratory observation, to be revisited when pan-genome data become available for both species.

The preceding analyses each related a single genomic or evolutionary feature to co-expression conservation separately. To identify which are independent predictors, we modelled conservation breadth (the per-gene number of stress–tissue comparisons with a conserved co-expressolog, 0–4) jointly against them using proportional-odds ordinal regression, complemented by linear regression and random-forest importance (Supplementary Figure S5). Once the co-expression-intrinsic properties (network degree and expression level) were held constant, only two features independently predicted conservation breadth: coding-sequence constraint (lower dN/dS; odds ratio 0.87 per standard deviation, P = 1 × 10⁻¹⁴) and shared segmental duplication (1.54-fold higher odds, P = 3 × 10⁻¹⁹), whereas presence–absence variation and lineage-specific duplication did not. Network degree itself carried an independent evolutionary signal with more highly connected genes being under stronger purifying selection (network degree versus dN/dS Spearman ρ = −0.11, P = 5.2 × 10⁻³⁵; median dN/dS declining across degree quartiles, 0.34, 0.33, 0.31 and 0.28). As such, network connectivity, coding-sequence constraint and conservation breadth vary together as facets of a single axis rather than as independent effects. The apparent dominance of network degree in the 1:1 model should be read with caution, as the 1:1 orthologue set has higher and more narrowly distributed expression than the full co-expression universe. The robust, model-independent findings are therefore the direction and significance of each predictor – degree positive, dN/dS negative, both independent, rather than their relative magnitudes. The same axis was recovered in the reciprocal Scots pine analysis: network degree was negatively correlated with cross-species dN/dS (Spearman ρ = −0.15, P = 1.9 × 10⁻⁵⁴), and degree and sequence constraint remained independent positive and negative predictors of conservation breadth (degree odds ratio 2.34, dN/dS odds ratio 0.85; both P < 1 × 10⁻¹⁸), with the expression range-restriction weaker than in Norway spruce, indicating the axis is not an artefact of subset composition.

## Discussion

Our central finding is that the breadth of cross-species co-expression conservation is coupled to coding-sequence constraint. Genes whose co-expression neighbourhoods are conserved most broadly between Norway spruce and Scots pine tend to be under stronger purifying selection, while the most regulatorily diverged are the least constrained. The gradient is modest in magnitude (Spearman ρ = −0.18, around 3% of variance in dN/dS) but directionally robust. These two species share a substantial, coordinated stress-regulatory response, with a large extent of shared differential expression at the orthogroup level and, beyond it, a further conserved regulatory backbone recovered by orthology-aware comparative co-expression that matched-timepoint expression comparison alone did not resolve. It was onto this shared backbone that the sequence-level signal of selection could then be mapped. The comparative co-expression approach is not specific to deeply diverged species, but this system is a particularly favourable window for coupling it to the selection signal. Cross-species co-expression retained across the ∼140-My *Picea–Pinus* split is strong evidence of functional constraint rather than shared ancestry, while the slow gymnosperm molecular clock keeps synonymous sites from saturating even at this depth, so dN/dS remains an interpretable measure of selection on the conserved core. Read together, comparative co-expression and molecular evolution define a single conservation–divergence axis, along which the conserved, network-central and constrained core can be distinguished from a heterogeneous lineage-specific periphery of reactive and structurally variable change. The sections below develop this axis, beginning with the organ- and species-specific structure of the response and moving to its evolutionary and genomic correlates.

### Organ identity structures the stress response

Across all datasets, organ identity was the dominant source of variance (PC1), with drought responses predominating in roots and cold responses in needles (Figure 1a–d). Under drought, roots showed a threshold-like transcriptional surge at the point of photosynthetic failure, followed by partial reversion after re-watering (Figure 1e), whereas needles mounted comparatively smaller responses. Under cold, needles responded earlier and more strongly than roots, most markedly in Scots pine. The limited overlap of individual DEGs between organs (Figure 2a–b), together with a cross-species orthogroup overlap that was substantial overall but accumulated only as each stress intensified (Figure 2c), underlines this extensive context specificity. The consistent among-sample clustering by stress and organ in the PCAs (Figure 1a–d) indicates that shared structure was present but this was not fully captured by DEG overlap.

### A conserved core of ABA, GA and growth control

Conserved drought-associated expression patterns, including enrichment of the conserved co-expression response for ABA-centred protective signalling (and, in Scots pine, stomatal-complex patterning genes), are congruent with ABA-driven stomatal closure that stabilises hydraulic function under declining water potential (Hasan et al. 2022; Albert et al. 2017; Postiglione and Muday 2020). Both species followed a shared, conservative water-use strategy, maintaining stomatal conductance until soil water became severely limiting before a late, sharp decline (this study; Haas et al. 2021; Meinzer et al. 2009); under a more acute deficit, needle death occurred in Norway spruce (Haas et al. 2021), consistent with hydraulic collapse as a recognised cause of drought-induced mortality in conifers (Choat et al. 2018; Arend et al. 2021). For cold, Ca²⁺/ROS signalling and the ICE–CBF–COR pathway are the canonical route to rapid protection via membrane and lipid remodelling and induction of cold-responsive genes. Although conifer ICE/CBF configurations differ from angiosperms and the canonical ICE1–CBF–COR module is not part of the conserved root cold response (Aro et al., in prep.), process-level conservation of the cascade, and of some ICE–CBF–COR components or close orthologues, is reported across plants (Hwarari et al. 2022; Vergara et al. 2022).

The cross-stress redeployment described above is most parsimoniously interpreted as conservation rather than convergence. The core ABA module (the PYL–PP2C–SnRK2 relay) and the GA–DELLA growth-repression module are ancient land-plant inventions. The ABA receptor and ABF-type transcription factor families arose with land colonisation, recruiting pre-existing PP2C phosphatases and SnRK kinases into a water-deficit network (Hauser et al. 2011; Komatsu et al. 2020); DELLA-mediated growth repression is conserved across land plants, its GA–GID1–DELLA perception module assembled in the vascular-plant ancestor (Hernández-García et al. 2021). Both modules thus predate the conifer–angiosperm divergence and the ∼140-My split of *Picea* and *Pinus* (Yeaman et al. 2014), and the GA–GID1–DELLA module remains demonstrably functional in conifers (Du et al. 2017). As reduced GA signalling with DELLA-mediated growth restraint is a broadly conserved response to cold, drought and other abiotic stresses (Achard et al. 2008; Colebrook et al. 2014), the shared GA-module enrichment under cold and drought in Norway spruce and Scots pine is better explained by independent redeployment of this conserved pathway than by convergent evolution of a new one. By the time the two genera diverged, the regulatory architecture coordinating growth arrest with stress tolerance was already in place. Lineage-specific divergence has, instead, concentrated at the periphery of this conserved core, among cold-specific and structurally variable genes, consistent with modification of an ancestral programme rather than its independent origin.

The conserved backbone recovered by comparative co-expression (Figure 3) carried a coherent hormonal logic. Among up-regulated drought-conserved co-expressologs, an ABA signature (Figure 4; Supplementary Table S4) is consistent with evidence from tree species that bZIP/ABF factors mediate ABA-induced stomatal closure and cytoskeletal remodelling controlling aperture (Yang et al. 2020; Huang et al. 2019). Under cold, prioritising protection over growth is expected to involve suppression of gibberellin (GA) activity, and GA-biosynthesis genes were correspondingly down-regulated under stress in both species (Supplementary Table S1), in line with the GA repression that accompanies cold acclimation in woody plants (Colebrook et al. 2014). In hybrid aspen (Populus tremula L. × P. tremuloides Michx.), low night temperature combined with inhibited GA biosynthesis overrides photoperiod to induce bud set and cold hardiness (Mølmann et al. 2005), underscoring the role of GA repression in cold acclimation of woody species (Olsen 2010). Persistent cell-cycle and growth down-regulation among drought co-expressologs, and cold-linked suppression of growth genes, are both consistent with hormone crosstalk in which ABA and reduced GA jointly stabilise physiology under stress (Hasan et al. 2022; Colebrook et al. 2014).

The conserved co-expression sets contained TF families canonically implicated in abiotic stress and development (Figure 3b): NAC, WRKY, MYB, AP2/ERF, bZIP/ABF, HSF, LBD and GLK. These plausibly form the conserved scaffolding of the stress response, on which lineage- and organ-specific fine-tuning of amplitude is superimposed; given the high cross-species correlation of expression trajectories, such tuning likely reflects subtle adjustments rather than wholesale differences.

### Distinguishing regulated signal from reactive change

Many organ- and stress-specific DEGs may reflect context-dependent rewiring or mis-regulation rather than novel stress-response pathways: cold and drought trigger genuine signalling cascades but also cause collateral membrane, oxidative and metabolic damage, so stress-induced expression mixes signalling-driven regulation with damage-associated response. The same causal-versus-consequence problem in disease genomics, where directionality-aware methods show many DEGs are consequences rather than drivers (Porcu et al. 2021), motivates separating the two, which matters directly for prioritising drivers as breeding or genome-engineering targets. Co-expression neighbourhoods coherent across conditions and species more likely represent core regulatory circuits than the condition-specific, fragmented signatures of damage, and network-preservation analyses similarly separate signalling hubs from genes tracking metabolic collapse (Zhang and Horvath 2005; Langfelder et al. 2011; Amrine et al. 2015). In our data, the disparity between limited within-species DEG overlap (Figure 2a–b) and strong cross-species neighbourhood conservation (Figure 3a–b) suggests many acute changes, particularly at photosynthetic failure under drought, are reactive, whereas conserved neighbourhoods are enriched for candidate core regulators (Furches et al. 2019; Curci et al. 2022). The condition-specific layer is itself heterogeneous, however, spanning genuine lineage-specific adaptation, including genes with population-genomic signatures of selection, as well as reactive change, so condition-specificity should not be equated with damage.

Comparative co-expression of wood formation across angiosperm and conifer trees, including Norway spruce and Scots pine, recovered more extensive conservation than seen here for stress (Rodríguez et al. 2026), consistent with tightly regulated developmental programmes resolving conserved regulation more cleanly than the less structured acute stress response. Biotic-defence categories (wounding, herbivory and oomycetes) were prominent among up-regulated genes in both species and stresses (Supplementary Table S4), consistent with abiotic–biotic cross-talk through shared hormone, ROS and MAPK modules and WRKY and NAC regulators (Atkinson and Urwin 2012), though this may be accentuated by transfer of GO annotations from angiosperm defence genes to conifer homologues and warrants caution.

### Conservation and divergence relative to angiosperm stress responses

Our characterisation of the conservation–divergence spectrum across five stress co-expression categories extends comparative analysis of abiotic stress regulatory-network conservation to a gymnosperm species pair. Kingdom-wide surveys show stress-response conservation decaying with phylogenetic distance yet remaining detectable across more than 700 million years of divergence (Koh et al. 2026), but the organisation of the stress response within conifers, and its coupling to coding-sequence and population-genomic variation, has remained relatively unexplored. These results can be set against patterns established in angiosperm systems. Cross-species comparisons in Arabidopsis, rice (*Oryza sativa* L.) and barley (*Hordeum vulgare* L.) have shown that 15–34 % of orthologous DEGs respond in qualitatively opposite directions between distantly related species, demonstrating that regulatory divergence is pervasive even where coding-sequence orthology is conserved (Hartmann et al. 2022). At closer phylogenetic distances, such as among hydroponic leafy-crop species, conserved gene regulatory networks are anchored by shared TF families (WRKY, AP2/ERF, GARP), with down-regulation of energetically demanding processes a broadly conserved feature of stress adaptation (Lee et al. 2025). The ∼1.9 % rate of diverged regulation we identified (genes stress-responsive in both species but with rewired co-expression neighbourhoods) measures a different quantity from the opposite-direction DEG rate of Hartmann et al. and the two are only loosely comparable. That these diverged-regulation genes also show elevated dN/dS indicates that regulatory rewiring is accompanied by relaxed coding-sequence constraint, a molecular signature of gene pairs that have partly escaped the co-regulatory architecture maintained in both species.

### Coding-sequence and structural signatures of conservation

The monotonic, albeit modest, decline of dN/dS with broader co-expression conservation places this regulatory gradient on a coding-sequence-evolution footing. Genes whose stress co-expression is conserved across both stresses and tissues tend to be under stronger purifying selection, whereas condition-restricted and non-conserved genes evolve faster on average. Regulatory-conservation status thus predicts coding-sequence constraint independently of network position. The association held in the joint model after accounting for network centrality and expression level, and is therefore not merely a by-product of conserved genes having higher network centrality. This gradient is consistent with the predominance of purifying selection (dN/dS below one) among forest-tree stress-response genes (Garosi et al. 2025). The strongest GO enrichment overall, RNA modification and organellar RNA processing among multi-tissue co-expressologs, suggests that tRNA and rRNA modification programmes constitute a general stress-regulatory infrastructure co-regulated across both stresses (Cai et al. 2025). Condition-specific functional signatures likewise align with prior conifer transcriptomics, with the drought-specific co-expressolog signatures echoing the transcriptional reprogramming reported in drought-stressed Scots pine needles (Zhou et al. 2024).

As shown above, the conservation–constraint coupling survived joint adjustment for network connectivity and expression and was independently reproduced by a within-species selection measure (pN/pS), so it is not simply a by-product of hub genes being both highly-connected and more strongly constrained (Mähler et al. 2017; Hämälä and Tiffin 2020). Two caveats bound this interpretation. First, dN/dS is defined only for the 1:1 orthologues within each category, and the divergent categories are dominated by genes without a 1:1 orthologue (for example, most not_coex stress-responsive genes lack one), so the gradient is anchored on the 1:1 minority of the divergent layer rather than on all of its genes; sequence constraint and genome architecture are therefore complementary descriptions of a single conservation–divergence spectrum rather than competing measures applied to one gene set. Second, that 1:1 set has higher and more narrowly distributed expression, which restricts the range over which network degree can vary and inflates its apparent weight relative to constraint in the joint model; the independent contribution of coding-sequence constraint is therefore best read from the genome-wide model and the pN/pS replication rather than from the 1:1 model alone.

Shared segmental duplicates were enriched in conserved drought circuits, whereas lineage-specific duplicates were enriched among non-conserved genes, suggesting gene duplication and structural variation are associated with regulatory divergence lying mostly outside the cross-species-conserved layer, with only a minority of independently duplicated genes occurring within shared regulatory architecture. This asymmetric partitioning after duplication parallels the loss of stress-responsive cis-regulatory elements driving rapid divergence among duplicate genes (Zou et al. 2009). Divergence in promoter transposable-element family composition between duplicate copies likewise accompanies their expression divergence, consistent with TE superfamilies supplying stress-type-specific regulatory motifs co-opted by transcription factors such as CBF/DREB and NAC (Deneweth et al. 2022). The *CHS3* case study (Figure 6) illustrates this at single-gene resolution. The current study has one notable limitation in that the population-genomic components of this spectrum (pN/pS, presence-absence variation, selection scans and the climate-GWAS overlap) were available only for Norway spruce (Ahlgren Kalman et al. 2025). These population-level signatures are thus established in Norway spruce and remain to be confirmed in Scots pine when comparable data become available. A further consideration is that the expression compendia of the two species were generated in separate experiments, so between-species batch effects cannot be excluded at the level of individual expression values. The conservation measure is, however, robust to this: cross-species co-expression is compared through within-species network neighbourhoods, each built independently within a single experiment, rather than through direct cross-experiment comparison of expression levels, so a systematic between-experiment offset shifts neither the co-expression topology within a species nor its conservation between them.

### A hierarchy of adaptation across the conservation–divergence axis

Adaptation is distributed across evolutionary timescales along the conservation–divergence axis, not confined to the conserved core (Figure 7). Deep, repeatedly co-opted machinery sits at the conserved end, under the strongest purifying selection and most central in the network, matching the high centrality and broad expression of orthogroups repeatedly recruited for local climate adaptation across plants (Whiting et al. 2024), whereas recent, lineage- and population-specific adaptation accumulates towards the faster-evolving, network-peripheral and structurally variable periphery. This axis yields an operational criterion for prioritising candidate regulators: genes that are simultaneously broadly conserved in cross-species co-expression, under strong purifying selection (low dN/dS) and central in the co-expression network. By recovering a conserved regulatory backbone that differential-expression overlap partially masks, orthology-aware comparative co-expression flags these candidates and separates them from the heterogeneous lineage-specific layer of condition-specific and structurally variable genes.

**Figure 7.**
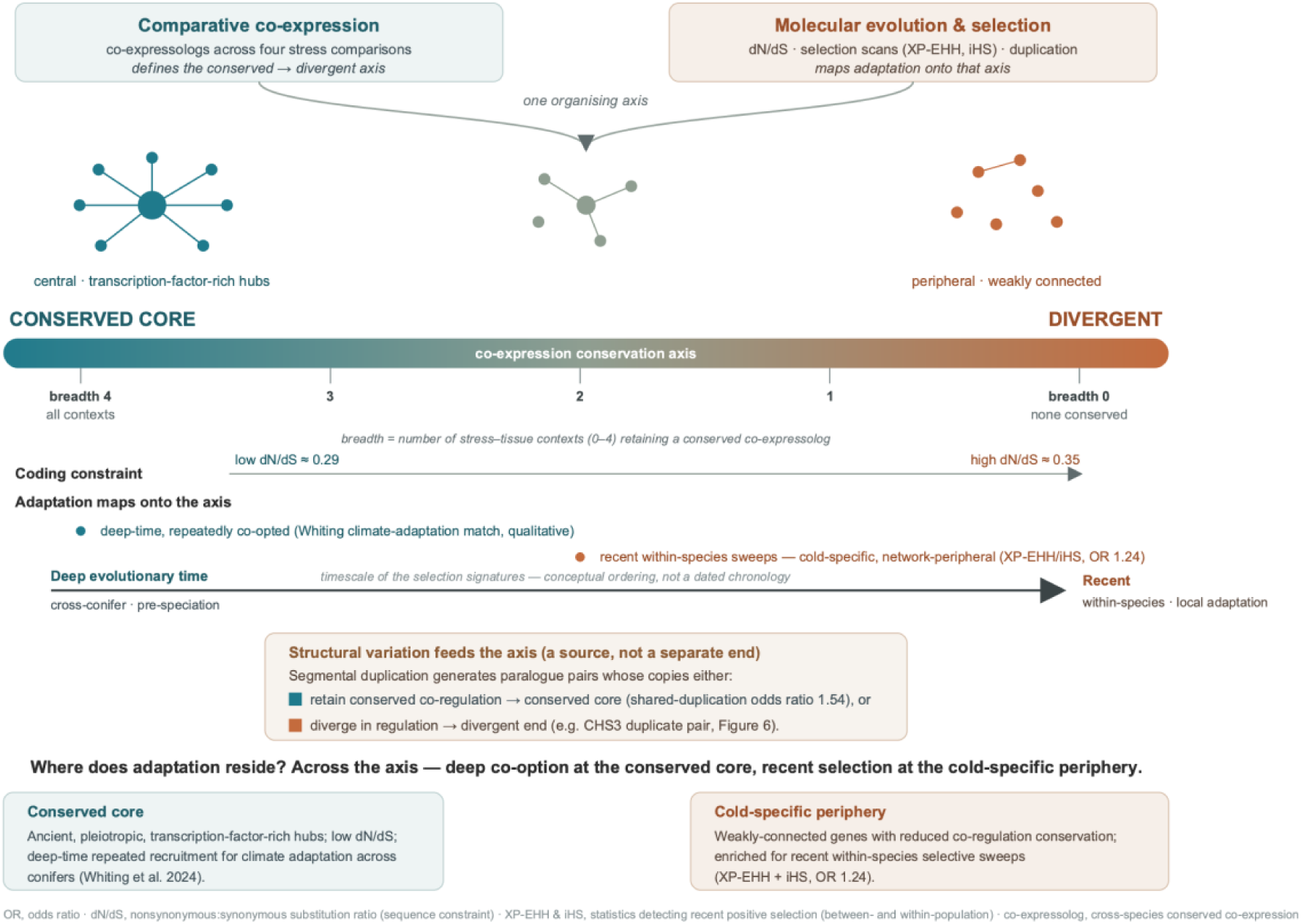
A single conservation–divergence axis integrates comparative co-expression with molecular evolution and selection. Cross-species co-expression conservation places each orthologous gene on a conserved-to-divergent axis (conservation breadth, the number of the four stress–tissue contexts in which a gene retains a conserved co-expressolog, 0–4); coding-sequence constraint (dN/dS ≈ 0.29 at the conserved core rising to ≈ 0.35 at the divergent end) and network position (central hubs to weakly connected periphery) vary along it. Two modes of adaptation map onto the axis: deep-time, repeatedly co-opted climate adaptation, which in other plants occupies central co-expression positions (Whiting et al. 2024) and recent within-species selective sweeps concentrated in cold-specific genes (odds ratio 1.24), which lie toward the network-peripheral, divergent end of the axis. Structural variation feeds the axis rather than forming a separate end: segmental duplication generates paralogue pairs whose copies either retain conserved co-regulation (shared-duplication odds ratio 1.54) or diverge in regulation (for example the *CHS3* pair, Figure 6). OR, odds ratio; dN/dS, non-synonymous:synonymous substitution ratio; XP-EHH and iHS, statistics detecting recent between- and within-population positive selection respectively.

## Materials and methods

### Plant material, experimental design and datasets

The Norway spruce drought stress data used here were previously published by Haas et al. (2021), and the Norway spruce cold stress data by Vergara et al. (2022). The 5 °C Scots pine root data were previously published in Aro et al. (in prep.). The 5 °C Scots pine needle data and the −5 °C Scots pine data (both organs) are newly presented in the current study. All datasets are available at the European Nucleotide Archive (ENA) under the umbrella accession PRJEB104157.

The drought experimental design of Haas et al. (2021) was replicated using three-year-old Scots pine seedlings of the Lilla Istad provenance (56° 30′ N), grown in pots of Hasselfors K-Jord (Hasselfors Garden, Örebro, Sweden) under long-day conditions (18 h light, 20 °C). Soil moisture was monitored gravimetrically as field capacity (FC), with seedlings at 80 % FC as well-watered controls. FC was reduced over five days, and needle and fine-root samples collected at 60 %, 40 % and 30 % FC. The 30 % FC condition, a mild, non-lethal drought stress, was maintained for seven days (30%7d) before severe drought was imposed by withholding water. Samples were taken at onset of visible photosynthetic dysfunction (Collapsed) and after two further days (C2d), after which seedlings were rehydrated to 80 % FC (Rehydrated). The stress trajectory was characterised physiologically via soil water content, shoot water potential, relative water content of shoots and roots, net photosynthesis and stomatal conductance (Supplementary Figure S1).

Cold stress was applied to Scots pine following the experimental design of Vergara et al. (2022). Two-year-old seedlings were acclimated to 18 °C, at which the first needle and root samples were taken. Seedlings were then transferred to 5 °C under 16 h light and sampled at 6 h, 24 h, 3 days and 10 days, after which they were transferred to −5 °C with additional samples taken at 6 h, 24 h, 3 days and 10 days. Five biological replicates were collected per timepoint and stored at −80 °C prior to RNA extraction.

### RNA extraction, sequencing and preprocessing

Samples were ground in liquid nitrogen and total RNA extracted following Chang et al. (1993), as modified by Street et al. (2006). Sequencing used an Illumina HiSeq 2500 platform with paired-end reads (2 × 126 bp). Raw reads were quality-assessed with FastQC (Andrews 2014) after each preprocessing step. rRNA was removed with SortMeRNA v1.9 (Kopylova et al. 2012) using default databases, adapters were removed and low-quality bases trimmed with Trimmomatic v0.39 (Bolger et al. 2014; ILLUMINACLIP:TruSeq3-PE.fa:2:30:10:1:TRUE SLIDINGWINDOW:5:30 MINLEN:38), and transcript abundance quantified with Salmon v0.11.3 (Patro et al. 2017) via selective alignment to reference genomes (*P. abies* v2 and *P. sylvestris* v1; Ahlgren Kalman et al. 2025). Counts were VST-normalised in DESeq2 v1.52.0 (Love et al. 2014) for quality assessment, and PCA (DESeq2 plotPCA, 500 most variable genes) of the first two components identified the major sources of variation within each dataset.

### Differential expression and functional enrichment

Differentially expressed genes (DEGs) were identified with DESeq2 (v1.52.0; Love et al. 2014), contrasting each stress condition against its control; genes with |log2 fold change| ≥ 2 and adjusted P-value ≤ 0.01 were retained. Heatmaps were generated with ComplexHeatmap v2.24.1. All R-based analyses ran in R 4.5 under Bioconductor 3.21 (with corresponding DESeq2, topGO, ComplexHeatmap and dependency versions). GO enrichment used topGO v2.60.1 with the weight01 algorithm (accounting for GO graph structure) and Fisher’s exact test (Alexa et al. 2006); co-expression category analyses were assessed against the ComPlEx expression universe (all genes retained in the comparative co-expression analysis; see below), again with weight01 and Fisher’s exact test. GO annotations came from the eggNOG-mapper and InterPro assignments of each gene; reported enriched terms were restricted to those in the Arabidopsis GO universe (org.At.tair.db), a reproducible plant-relevance filter rather than annotation transfer from Arabidopsis.

For the cross-species orthogroup-level comparison (Figure 2c), differentially expressed genes from each species were mapped to orthogroups, and the analysis was restricted to the 11,943 orthogroups that contain at least one gene from both Norway spruce and Scots pine. Differentially expressed genes belonging to single-species orthogroups (1,027 Norway spruce and 1,461 Scots pine genes, 8.5% and 10.9% of the differentially expressed genes of each species) have no possible counterpart in the other species and were therefore excluded from this comparison. Overlap percentages are expressed relative to the species with the smaller differentially expressed orthogroup set at each sampling point.

### Orthology and identification of co-expressologs

Gene families (orthogroups) were identified using OrthoFinder (Emms and Kelly 2019), using results from Ahlgren Kalman et al. (2025). Conserved co-expression relationships were identified using Comparative analysis of Plant co-Expression (ComPlEx; Netotea et al. 2014), following the comparative co-expression approach of Rodríguez et al. (2026). For each species and condition, a co-expression network was constructed from Pearson correlations of the filtered expression matrices, mutual ranking applied to improve robustness, and networks trimmed to retain the strongest 3% of possible edges (density = 0.03). For each orthologue pair, a bidirectional hypergeometric test assessed whether co-expression neighbourhoods overlapped significantly between species, with Benjamini–Hochberg correction for multiple testing; pairs significant in both directions were termed co-expressologs. As only two species are compared here, conserved co-expression is assessed directly between orthologue pairs. Comparisons were conducted between equivalent individual networks (for example, Norway spruce drought needle against Scots pine drought needle), and co-expressologs were retained at a bidirectional adjusted P-value (the larger of the two reciprocal species P-values, denoted MaxpVal) below 0.01, so that a retained co-expressolog is significant in both directions. ComPlEx was applied using a Python re-implementation developed for this study, which reproduces the original R implementation (https://gitlab.com/hvidsten-lab/rcomplex) exactly, producing identical co-expressolog calls on a matched validation subset (100% concordance) while running substantially faster (https://github.com/natstreet/ComPlEx_python). Network degree was computed for each gene as its mean co-expression degree across the four Norway spruce ComPlEx stress networks (cold and drought, needles and roots), averaged over the networks in which the gene was present.

### Classification of cross-species co-expression conservation

Genes were assigned, through their orthogroups, to five mutually exclusive categories based on cross-species co-expression of their co-expressolog pairs across the four stress–tissue comparisons (cold and drought, needles and roots). For each orthogroup, pairs were scored by tissue using the NegLog10 co-expression evidence (−log10 MaxpVal per comparison; comparisons lacking a significant co-expressolog contributed zero), summed across relevant comparisons. Conserved orthogroups had significant pairs (MaxpVal < 0.01) in all four comparisons with a summed score ≥ 10 across all four tissues. Cold-specific orthogroups had pairs in the cold comparisons (needle and/or root) but none in either drought comparison (ColdSum ≥ 5, DroughtSum = 0), and drought-specific orthogroups were defined analogously. Multi-tissue orthogroups had pairs in two or three comparisons spanning both stresses that did not meet the single-stress criteria (summed score ≥ 5), named for recurring across multiple stress–tissue conditions rather than a single stress. Not co-expressed (not_coex) orthogroups had no significant cross-species co-expression in any comparison. Category assignment was hierarchical: orthogroups assigned a higher-priority category were excluded from lower ones. Species-specific stress responses were then characterised among the stress-responsive not_coex genes: Norway spruce genes differentially expressed in at least one stress comparison and assigned to not_coex. Each was classified by the reason it lacked a conserved co-expressolog into one of four mechanisms: no 1:1 orthologue in the other species (whether lineage-specific or in a non-1:1, expanded gene family), a 1:1 orthologue not itself stress-responsive, diverged regulation (both species stress-responsive but with divergent co-expression neighbourhoods), or a 1:1 orthologue not expressed under stress (Supplementary Figure S3a). This stress-responsive not_coex set is distinct from, and larger than, the subset of not_coex genes carrying a 1:1 orthologue that enter the dN/dS backbone, because most stress-responsive not_coex genes have no 1:1 orthologue. The equivalent classification was performed for Scots pine.

### Conserved co-expressolog expression dynamics

For each stress, conserved co-expressolog orthogroups were defined as those differentially expressed in both species and showing strong conserved cross-species co-expression in the corresponding tissues (summed −log10 co-expression evidence ≥ 60 for drought and ≥ 40 for cold); where an orthogroup contained more than one qualifying pair, the pair with the highest summed co-expression evidence was retained as the representative. Expression-profile conservation was quantified per orthogroup and tissue as the Pearson correlation between Norway spruce and Scots pine median expression trajectories across matched physiological stages of the stress time-course and compared with a null distribution obtained by shuffling the Scots pine partner among orthogroups (fixed random seed, set.seed(42)). Each orthogroup was assigned an up- or down-regulated direction from the sign of the mean late-stress minus baseline expression in Norway spruce, averaged across tissues. GO enrichment was computed separately for the up- and down-regulated sets as described above.

### dN/dS estimation

Rates of non-synonymous (dN) and synonymous (dS) substitution were estimated for 13,523 cross-species orthologue pairs that are 1:1 between Norway spruce and Scots pine, defined as OrthoFinder pairwise orthogroups containing exactly one gene from each species. For each orthologue pair, coding sequences were translated and aligned at the protein level using a global pairwise alignment (BioPython PairwiseAligner, BLOSUM62 substitution matrix; gap-open = −11, gap-extend = −1) and back-translated to a codon alignment. dN and dS were then estimated under the Yang and Nielsen (2000) maximum-likelihood model using the yn00 program of PAML v4.10.10 (icode = 0, weighting = 0, commonf3x4 = 0). Pairs were excluded if either translated protein was shorter than 20 residues or contained an internal stop codon; in downstream analyses, pairs with dS outside the interval (0, 5) or dN/dS > 10 were additionally excluded as unreliable estimates, leaving 13,519 pairs with usable dN/dS.

### Segmental duplication identification

Segmental duplicate (SD) genes were classified as duplicated independently in both Norway spruce and Scots pine (shared_SD) or only in the Norway spruce lineage (spruce_only_SD). SD gene lists for both species come from Ahlgren Kalman et al. (2025), available in the associated SciLife FigShare repository. Briefly, duplicate gene blocks were identified as genomic regions containing two or more genes/pseudogenes with at least two copies in the focal species, using DBSCAN cluster detection and syntenic block analysis implemented in R (scripts at https://github.com/tallgran/conifer-JasonHill). SD genes were cross-referenced with N10 hierarchical orthogroup (HOG) membership: an HOG was classified as shared_SD if SD-block genes occurred in both Norway spruce and Scots pine, indicating independent duplication of the same ancestral gene in both lineages, and as spruce_only_SD if SD-block members occurred only in Norway spruce. The same classification was applied reciprocally in Scots pine, using the Scots pine segmental-duplication calls (Ahlgren Kalman et al. 2025; same DBSCAN and syntenic-block method) and the Scots pine co-expression category assignments, to define a pine_only_SD class (SD-block members occurring only in Scots pine) and test its enrichment across the Scots pine co-expression categories. The population-genomic measures integrated later (presence–absence variation, pN/pS and selection scans) remain available only for Norway spruce, as no equivalent Scots pine population resequencing framework exists. Independent rather than ancestral origin of shared_SD duplicates was confirmed by Ks dating: median within-Norway spruce Ks of SD pairs was approximately 40–53% of cross-species Ks (calibrated to the ∼140 Mya *Picea–Pinus* divergence), placing most duplications after species separation.

### Presence–absence variation

Presence–absence variation (PAV) was assessed for genes across a standing population panel of Norway spruce using PAV calls from Ahlgren Kalman et al. (2025) and are available in the associated SciLife FigShare repository. Briefly, a gene was scored as absent in an individual if it showed no detectable read coverage across the gene body in whole-genome resequencing data from the Norway spruce standing population panel described in Ahlgren Kalman et al. (2025) Genome-wide XP-EHH and iHS selection scans and climate-variable genome wide association study (GWAS) results were likewise obtained from Ahlgren Kalman et al. (2025). Genes were classified as carrying both selection signals on this basis (n = 2,712). For these genes we compared, against no-selection background genes, two properties over the within-species co-expression network-node set: co-expressolog degree (the count of conserved cross-species co-expressolog partners) and expression level (the mean DESeq2 baseMean of each gene across the four Norway spruce stress experiments; cold and drought, needles and roots).

### Comparison with a developmental co-expression dataset

To assess whether the genomic signatures associated with stress co-expression conservation are specific to stress or reflect more general properties of conserved regulation, we obtained the wood-formation (developmental) co-expression data for Norway spruce and Scots pine from Rodríguez et al. (2026) and applied the same dN/dS and presence–absence-variation overlays used for the stress co-expression categories above.

### Transcription-factor binding motifs in the *CHS3* case study

Transcription factors were annotated in the Norway spruce gene models using PlantTFDB (Jin et al., 2017), yielding 1,954 annotated TFs. The transcription-factor composition in Figure 3b summarises orthogroups whose co-expression is conserved across both species. As such, the Norway spruce member was used as the annotation reference for each conserved pair, each conserved co-expressolog orthogroup containing a Norway spruce gene whose family assignment represents the shared orthogroup, so a separate Scots pine annotation was not required. For the *CHS3* segmental-duplicate case study (Figure 6), transcription-factor binding motifs in the 2 kb region upstream of the transcription start site of each copy were predicted with FIMO (version 5.5.9, MEME Suite; Grant et al. 2011; P ≤ 1 × 10⁻⁴) against the PlantTFDB motif database, and the sets of significant motif families in the 2 kb upstream region of the two gene copies were compared. The Norway spruce chalcone synthase gene family (orthogroup OG0000177; 20 *Picea abies* members) was aligned at the protein level with MAFFT (--auto) and a maximum-likelihood phylogeny inferred with FastTree under the LG substitution model, retaining SH-like local branch-support values (Supplementary Figure S4). Pfam domain assignments for the *CHS3* copies (chalcone-synthase N- and C-terminal domains, PF00195 and PF02797) were taken from the InterPro Pfam annotation of the Norway spruce gene models, with the same domains independently assigned by eggNOG-mapper.

### Integrative model of conservation

To identify genomic and evolutionary features independently predicting co-expression conservation, conservation breadth (0–4) was modelled by proportional-odds ordinal regression against standardised predictors. dN/dS is defined only for 1:1 orthologues, whereas segmental-duplication and presence–absence-variation status apply to duplicate genes, so two complementary models were fitted: a genome-wide model across all co-expression-universe genes (predictors: network degree, mean expression, shared and lineage-specific segmental-duplication status, and presence–absence variation) and a 1:1-orthologue model (predictors: dN/dS, promoter transposable-element divergence, network degree and mean expression). Promoter transposable-element divergence was quantified for 1:1 orthologues as the Jaccard distance between transposable-element families in the 2 kb promoter region of the Norway spruce and Scots pine copies. The promoter was defined strand-aware relative to the representative (longest-CDS) transcript: the 2 kb interval immediately upstream of the transcription start site, taken as the gene start for plus-strand genes and the gene end for minus-strand genes. Transposable-element families were taken from the RepeatMasker repeat annotations of each genome (Picab02 and Pinsy01; family-level assignments). Continuous predictors (network degree, mean expression, dN/dS and promoter transposable-element divergence) were standardised to zero mean and unit variance (z-scores). The proportional-odds assumption was tested with the Brant test, and ordinal estimates were checked against standardised linear regression (95% confidence intervals) and random-forest permutation importance. To avoid dependence on any single random seed, permutation importance was computed as the mean over 100 random forests (1,000 trees each, fixed seeds 1 to 100), with the run-to-run standard deviation reported as a measure of stability. To confirm the conservation gradient was not an artefact of classification thresholds, category assignment was repeated across a range of co-expressolog evidence cut-offs (summed −log10 MaxpVal of 10–30 for the conserved category and 5–15 for the stress-specific and multi-tissue categories); the monotonic dN/dS ordering, its significance and the segmental-duplicate enrichment pattern were preserved throughout (Supplementary Table S7). This sensitivity analysis varied the co-expressolog evidence thresholds used to assign categories; the underlying network density (strongest 3% of edges) followed the value justified by saturation analysis in Rodríguez et al. (2026), at which the recovered co-expressolog set stabilises.

### Data availability

Raw RNA-seq reads from all experiments are deposited in the European Nucleotide Archive (ENA) under accession PRJEB104157, the umbrella accession that collates all datasets used in this study. The raw reads from the previously published Norway spruce datasets are accessible under the accessions given in Haas et al. (2021) and Vergara et al. (2022). Processed expression matrices, DESeq2 dataset objects (dds_objects.tar.gz), co-expression network outputs, genome annotation files, and key intermediate analysis files are archived on the SciLifeLab FigShare repository (doi: 10.17044/scilifelab.32593647). Genomic data for Norway spruce and Scots pine used in the orthology and segmental duplication analyses are available from ENA under accessions PRJEB69221 and PRJEB77112, respectively (Ahlgren Kalman et al. 2025). Population-level presence–absence variation data and segmental duplication gene lists are available from the SciLifeLab FigShare repository associated with Ahlgren Kalman et al. (2025) (doi: 10.17044/scilifelab.28737623).

### Code availability

Scripts used to perform the analyses presented here are available in the git repository at https://github.com/natstreet/conifer-stress-comparative-genomics (DOI: 10.5281/zenodo.21628206). The original R implementation of ComPlEx is available at https://gitlab.com/hvidsten-lab/rcomplex, and the Python re-implementation used to generate the co-expressolog results at https://github.com/natstreet/ComPlEx_python (DOI: 10.5281/zenodo.21629336).

Declaration of AI use: the Python re-implementation of the ComPlEx co-expressolog algorithm (Netotea et al. 2014) was developed with the assistance of a large-language-model coding assistant (Claude, Anthropic; Claude Opus 4 series, 2026). The generated code was validated against the original R implementation, yielding numerically equivalent co-expressolog calls (validation scripts are included in the repository), and the authors reviewed all AI-assisted code and analyses and take full responsibility for the content and accuracy of the manuscript.

## Supplementary Figures

**Supplementary Figure S1.**
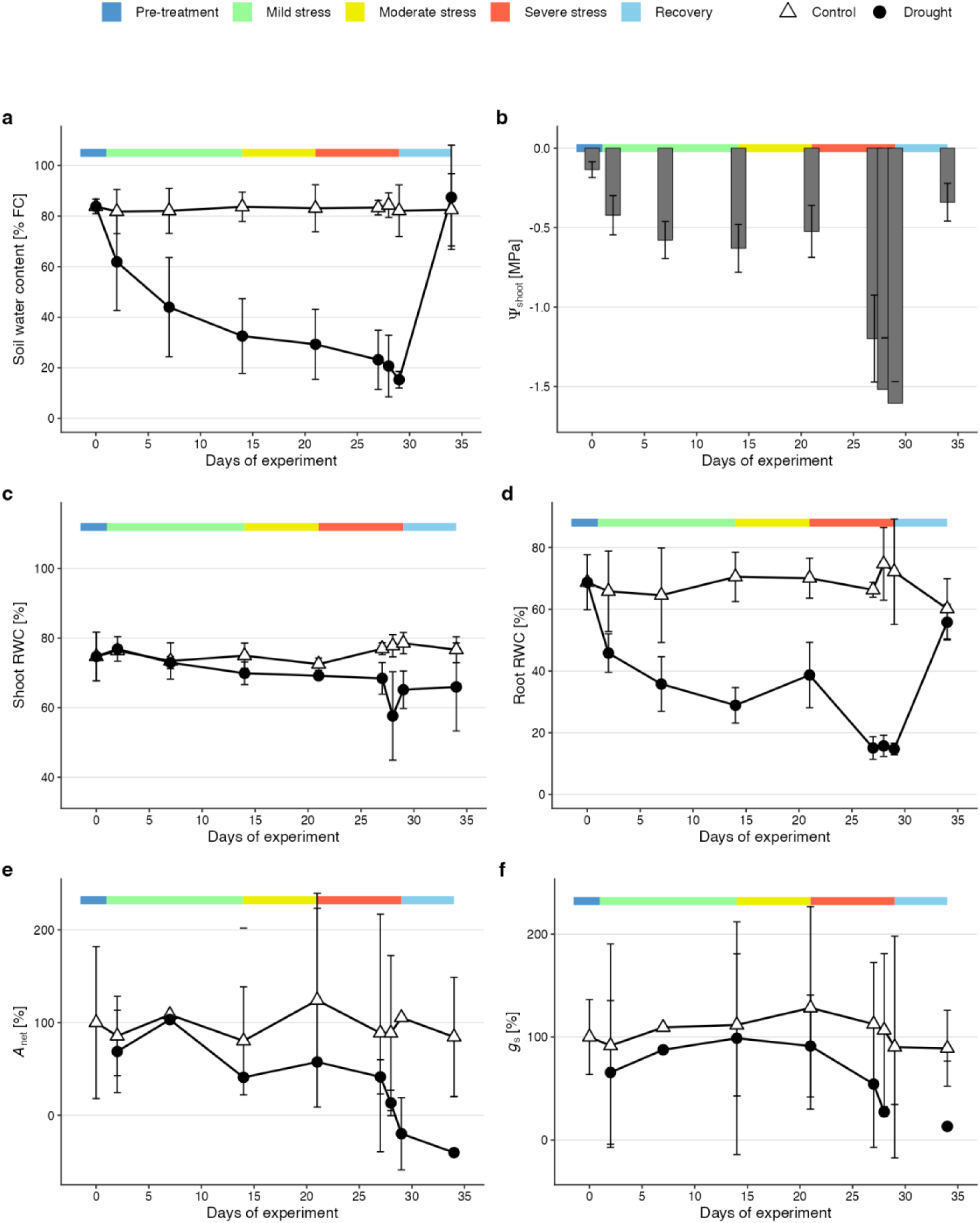
Physiological characterisation of the Scots pine drought experiment. (a) Soil water content as percentage of field capacity (% FC) in control and drought-treated plants throughout the 34-day experiment. (b) Shoot water potential (Ψshoot, MPa) in drought plants; well-watered controls maintained ∞ −0.25 MPa throughout. (c) Shoot relative water content (RWC, %). (d) Root RWC (%). (e) Net photosynthesis (Amax) and (f) stomatal conductance (gs) expressed as a percentage of the Day 0 pre-treatment reference value; values below zero indicate net CO₂ efflux. Points/bars represent means ± 95% confidence intervals (n = 5 per group per timepoint; n = 3 for gas exchange measurements). Coloured bars at the top of each panel indicate experimental phases (blue, pre-treatment; green, mild stress, FC 60–30%; yellow, moderate stress, FC ∼30%; red, severe stress and collapse, FC 22–15%; light blue, recovery, FC ∼80%).

**Supplementary Figure S2.**
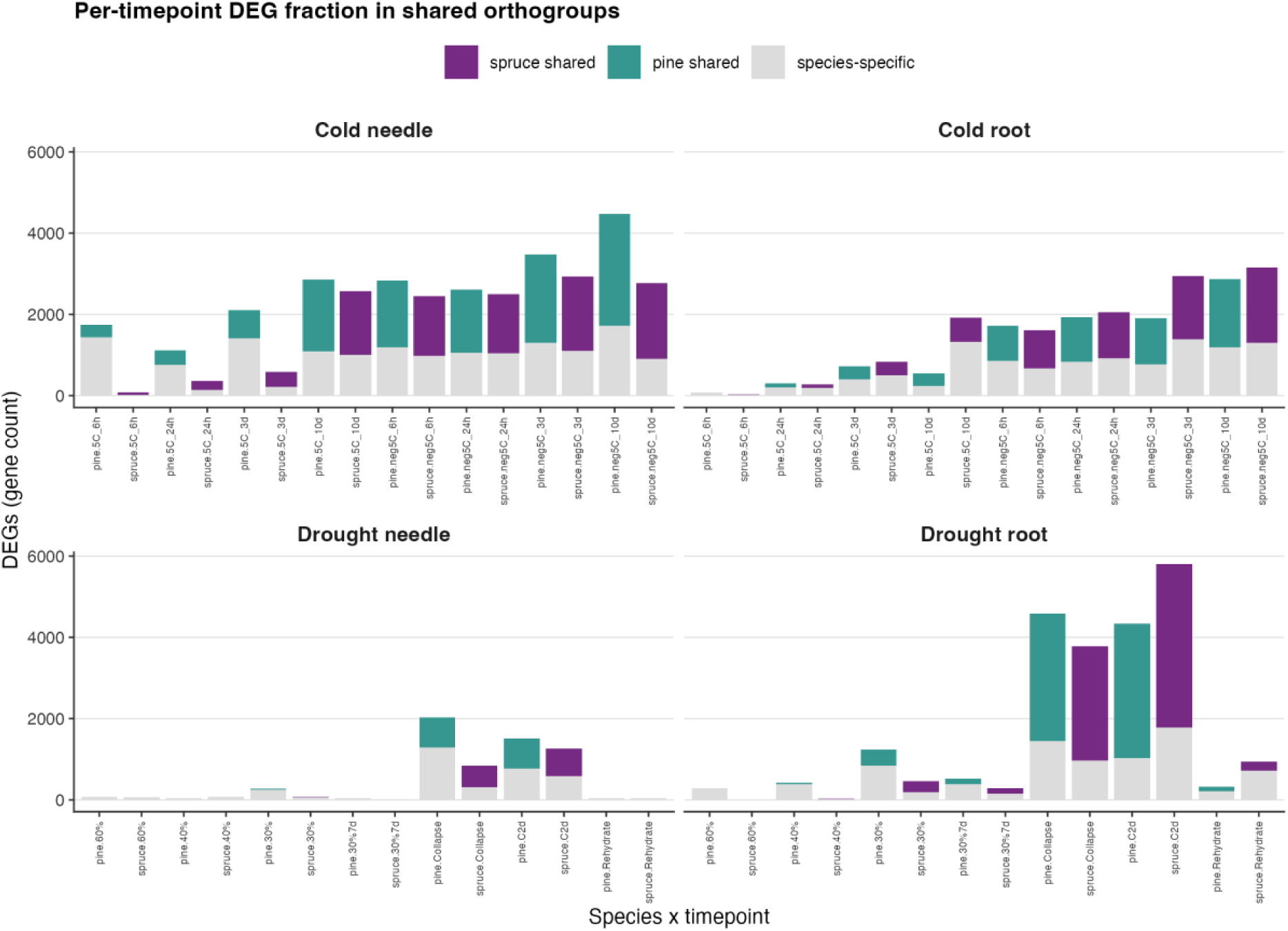
Cross-species orthogroup-level differentially expressed gene overlap resolved by sampling point. For each stress–tissue series, stacked bars show the number of differentially expressed genes (DEGs) at each sampling point that fall in orthogroups shared with the other species (coloured: Norway spruce shared, purple; Scots pine shared, teal) versus those in species-specific orthogroups or without a cross-species counterpart (grey). Panels are cold needle, cold root, drought needle and drought root; within each panel, paired bars give the two species at matched sampling points. The shared fraction is small at the earliest, mildest timepoints and rises steeply as stress severity increases, indicating that the substantial orthogroup-level overlap summarised in Figure 2c emerges progressively rather than being present uniformly across the stress progression.

**Supplementary Figure S3.**
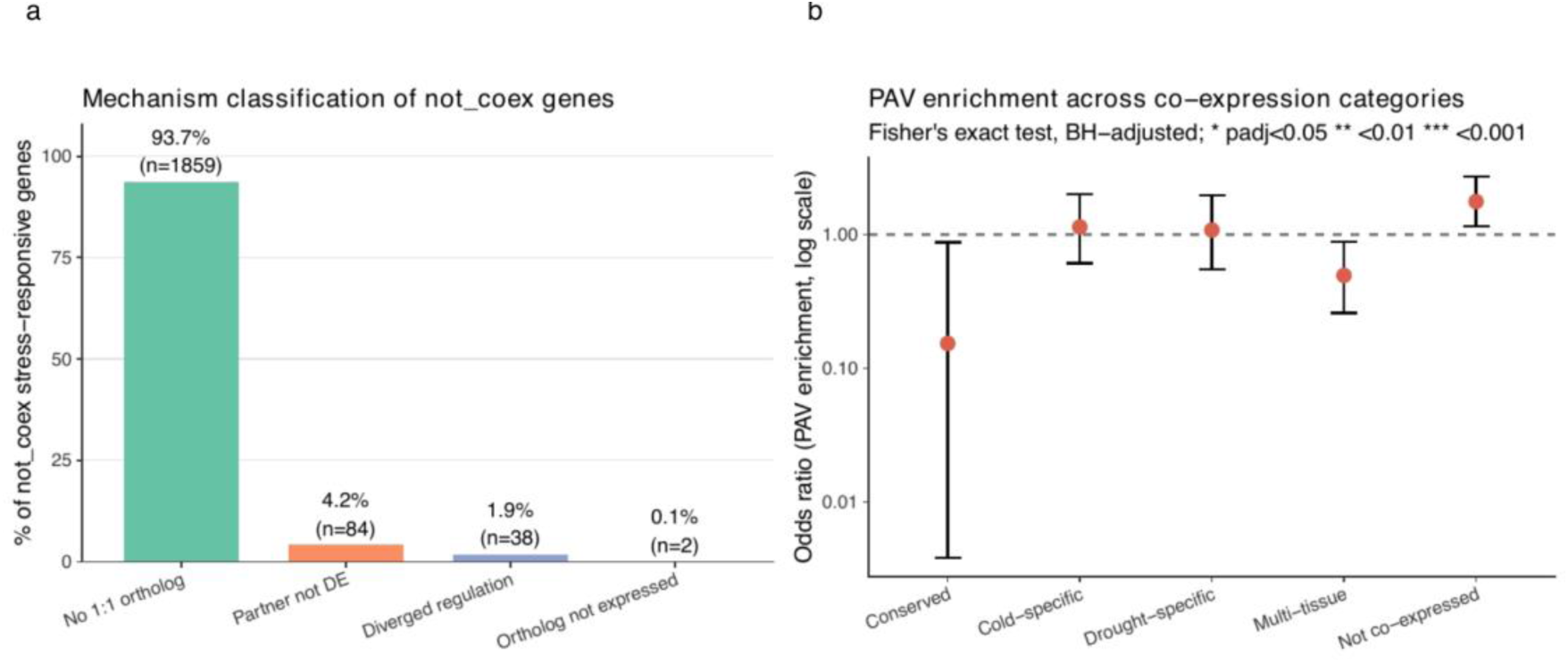
Mechanism classification and population-genomic features of not co-expressed stress-responsive genes. (a) Proportion of genes that are stress-responsive in at least one species but classified as not co-expressed (not_coex), across mechanism categories: no 1:1 orthologue; partner not DE; diverged regulation; orthologue not expressed (percentages and gene counts shown). (b) Odds ratio for enrichment of presence–absence-variable (PAV) genes from the *Picea abies* standing population across co-expression conservation categories (Fisher’s exact test, Benjamini–Hochberg correction). Points show the odds ratio; the dashed line marks OR = 1 (no enrichment). No category showed enrichment surviving Benjamini–Hochberg correction (all Padj > 0.05).

**Supplementary Figure S4.**
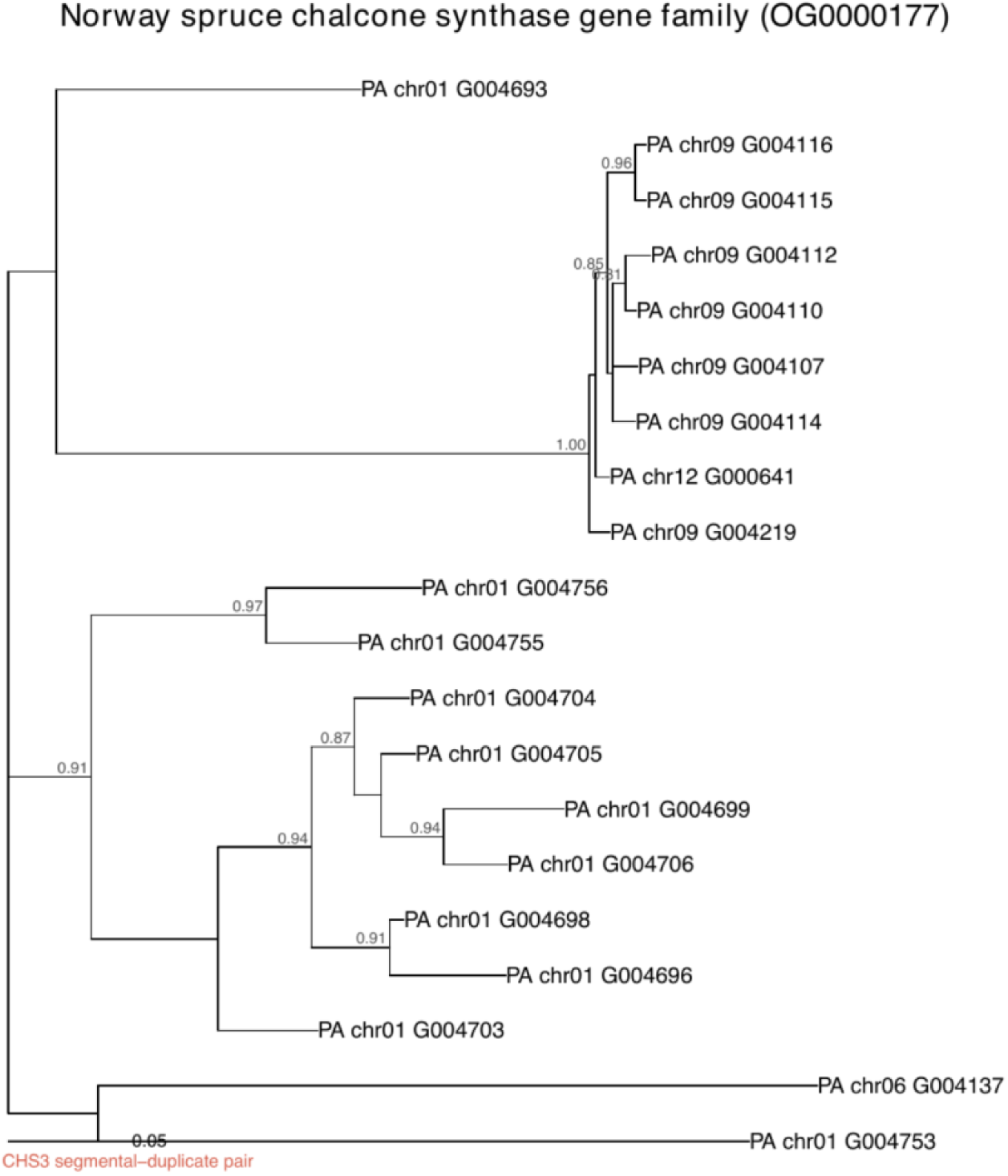
Maximum-likelihood phylogeny of the Norway spruce chalcone synthase gene family. Protein sequences of the 20 *Picea abies* members of the chalcone synthase (CHS) orthogroup (OG0000177) were aligned with MAFFT and a maximum-likelihood tree inferred with FastTree (LG model); SH-like branch-support values ≥ 0.8 are shown and scale is in substitutions per site. The segmental-duplicate *CHS3* pair (PA_chr09_G004115 and PA_chr09_G004116) forms a strongly supported sister pair (support 0.96) nested within the chromosome-9 chalcone synthase tandem array, consistent with a recent duplication. Both genes carry the diagnostic Chal_sti_synt_N (PF00195) and Chal_sti_synt_C (PF02797) Pfam domains.

**Supplementary Figure S5.**
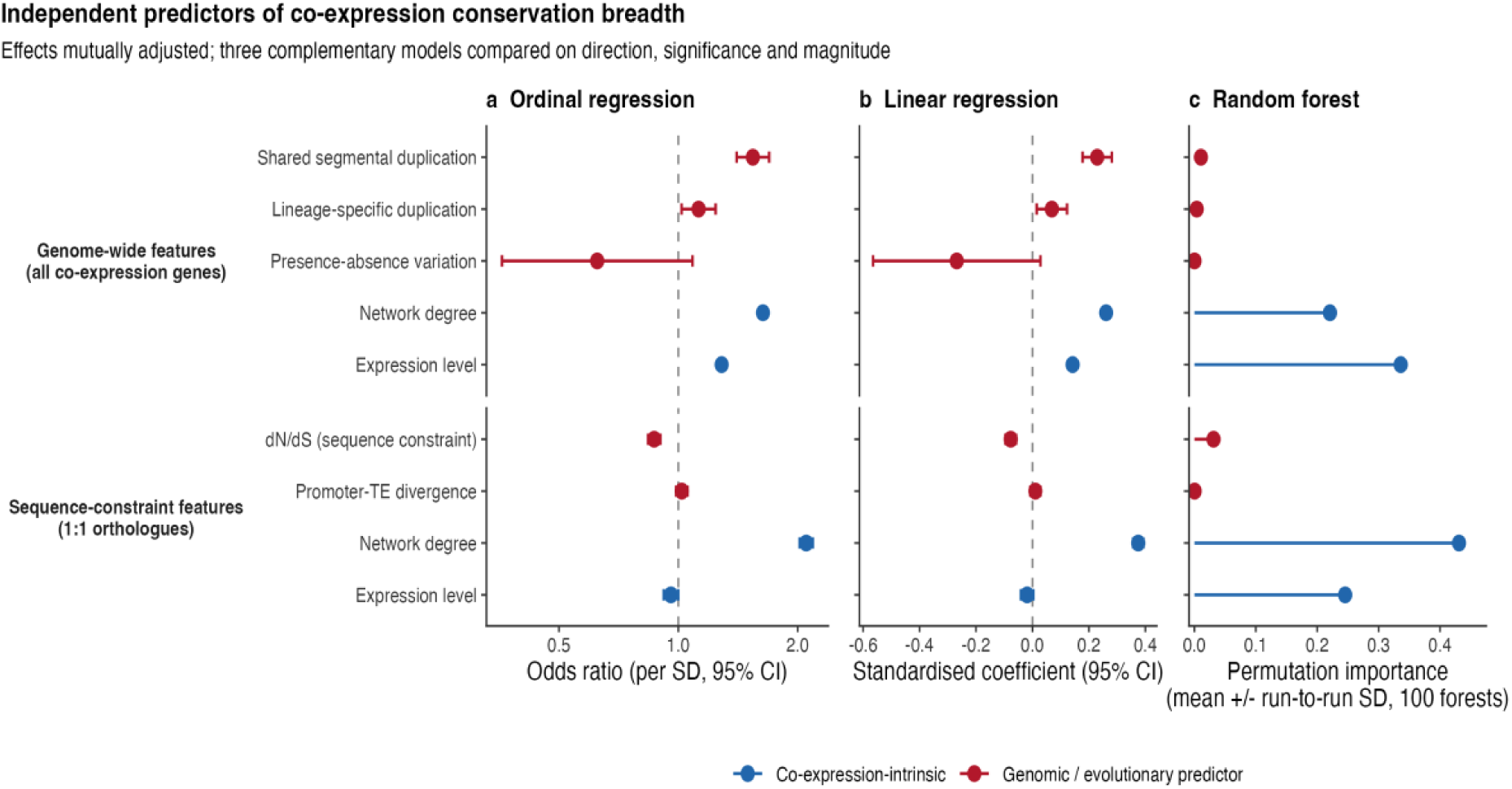
Genomic and evolutionary predictors of co-expression conservation. Effects on conservation breadth (0–4) estimated by three complementary methods, each from two complementary models: genome-wide features across all co-expression genes and sequence-constraint features across 1:1 orthologues. (a) Standardised effects as odds ratios with 95% confidence intervals from proportional-odds ordinal regression. (b) Standardised linear-regression coefficients with 95% confidence intervals. (c) Random-forest permutation importance, shown as the mean over 100 forests (1,000 trees each, fixed seeds 1 to 100); error bars are the run-to-run standard deviation, not a confidence interval. Effects are mutually adjusted within each model, and features share a common ordering across panels. Red, significant genomic/evolutionary predictors; blue, co-expression-intrinsic properties (network degree, expression level); grey, not significant. The three methods agree in direction and significance: dN/dS (negative) and shared segmental duplication (positive) independently predict conservation beyond the co-expression-intrinsic properties, while presence–absence variation and promoter transposable-element divergence do not. Network degree and expression level dominate in magnitude, consistent with the caution noted in the main text against over-interpreting the relative sizes of effects.

## Notes

https://doi.org/10.17044/scilifelab.32593647

### Competing Interest Statement

The authors have declared no competing interest.

### Funding

- Trees and Crops for the Future

